# Genome streamlining: effect of mutation rate and population size on genome size reduction

**DOI:** 10.1101/2024.03.14.584996

**Authors:** Juliette Luiselli, Jonathan Rouzaud-Cornabas, Nicolas Lartillot, Guillaume Beslon

## Abstract

Genome streamlining, *i.e*. genome size reduction, is observed in bacteria with very different life traits, including endosymbiotic bacteria and several marine bacteria, raising the question of its evolutionary origin. None of the hypotheses proposed in the literature is firmly established, mainly due to the many confounding factors related to the diverse habitats of species with streamlined genomes. Computational models may help overcome these difficulties and rigorously test hypotheses. In this work, we used Aevol, a platform designed to study the evolution of genome architecture, to test two main hypotheses: that an increase in population size (*N*) or mutation rate (*µ*) could cause genome reduction. In our experiments, both conditions lead to streamlining, but with very different resulting genome structures. Under increased population sizes, genomes loose a significant fraction of non-coding sequences, but maintain their coding size, resulting in densely packed genomes (akin to streamlined marine bacteria genomes). By contrast, under increased mutation rate, genomes loose both coding and non-coding sequences (akin to endosymbiotic bacteria genomes). Hence, both factors lead to an overall reduction in genome size, but the coding density of the genome appears to be determined by *N × µ*. Thus, a broad range of genome size and density can be achieved by different combinations of *N* and *µ*. Our results suggest that genome size and coding density are determined by the interplay between selection for phenotypic adaptation and selection for robustness.

**Significance statement:** Many bacterial species show reduced genomes. However, the diversity of these species and of their life traits makes it difficult to identify the mechanisms that led to this reduction. Indeed, no unifying hypothesis accounts for the whole diversity of genome size reduction. Here, we used simulations to systematically explore the effect of population size and mutation rate on genome size. We show that the interaction between these two factors tightly determine the size, but also the density of genomes, making it possible to account for the whole diversity of reduced genomes by acting on these two parameters only. Our results suggest a theoretical model in which genome reduction is driven by a robustness/fitness trade off.

## 1 Introduction

Genome size was one of the first studied genome characteristics (Leth Bak et al., 1969; Bachmann, 1972), yet its dynamic and causal factors are still poorly understood. Genome size is hugely variable across life: from less than 10^4^ base pairs (bp) for viruses (Gago et al., 2009), to more than 10^11^ bp for some plants (Pellicier et al., 2010). It does not correlate reliably with the number of genes or other variables throughout the different branches of life (Barow and Meister, 2002; Westoby et al., 2021).

The observed range of genome sizes is largely reduced when studying only bacterial organisms (Westoby et al., 2021), ranging from 10^5^ bp for intracellular endosymbiotic bacteria (Chong et al., 2019) to 10^7^ bp for some myxobacteria (Schneiker et al., 2007). Bacterial genomes are mostly dense, and within this domain of life, genome size is loosely correlated with the number of coding genes (Konstantinidis and Tiedje, 2004; Almpanis et al., 2018). However, the precise determinants of bacterial genome size are still unknown, as it is still impossible to accurately predict the total genome size from the number of coding genes or from other genomic characteristics (Petrov, 2001; Barow and Meister, 2002; Choi et al., 2020). Part of the determinants of genome size are likely to be highly lineage-specific and linked to the ecological or evolutionary history of the lineages (Martinez-Gutierrez and Aylward, 2022). Nevertheless, it has been argued that at least a part of the observed variation may be due to universal mechanisms, linked to population genetics and molecular evolutionary processes (Lynch and Conery, 2003; Lynch, 2007). In particular, it has been suggested that population genetics mechanisms could explain the reductive evolution observed in several bacterial strains (Lynch, 2006). However, among the shortest bacterial genomes, one can find two types of bacteria which have very different ecological environments and evolutionary history: endosymbionts such as *Buchnera aphidicola* (Moran and Mira, 2001) *and free-living marine bacteria such as Prochlorococcus marinus* (Dufresne et al., 2005) *or Pelagibacter ubique* (Giovannoni et al., 2005). *Strikingly, both types of bacteria lie at the two extremes of bacterial population sizes, questioning the mechanisms that led to genome reduction (Batut et al., 2014; Martínez-Cano et al., 2015; Wernegreen, 2015)*.

*Buchnera aphidicola*, and endosymbionts more generally, are characterized by very small effective population sizes (*N*_e_) and high mutation rates (*µ*). Endosymbiosis also generally entails the introduction to a new stable environment and very close interactions with the host (Moran, 1996; Mira and Moran, 2002). These many complex factors result in decaying genomes, smaller in total size and with less coding genes than those of average bacteria (Heddi et al., 1998). Endosymbionts have typically lost both coding and non-coding genomic content (Moran and Mira, 2001; Wernegreen, 2002), maintaining a coding fraction on the order of 85% (van Ham et al., 2003), which is quite typical for bacteria (Kuo et al., 2009).

In sharp contrast, free-living marine bacteria such as *Prochlorococcus marinus* or *Pelagibacter ubique* also have reduced genomes (Giovannoni et al., 2005; Batut et al., 2014), but are believed to have very large effective population sizes (Marais et al., 2008; Flombaum et al., 2013; Giovannoni et al., 2014), although that is an ongoing debate (Chen et al., 2022; Filatov and Kirkpatrick, 2024). Noticeably, in their case, genome size reduction is primarily contributed by the loss of non-coding sequences rather than coding sequences (Giovannoni et al., 2005; Batut et al., 2014). This phenomenon is called streamlining and could indicate a very effective selection (Wolf and Koonin, 2013; Giovannoni et al., 2014). Many hypotheses have been proposed to account for genome size reduction and the associated changes in genome architecture in such free-living organisms: adaptation to a nutrient poor environment or to other abiotic factors, the Black Queen hypothesis, or high mutation rates (Koskiniemi et al., 2012; Morris et al., 2012; Batut et al., 2014; Ngugi et al., 2023).

Both endosymbionts and free-living marine bacteria thus show a marked reduction in genome size, linked to an increase in mutation rate (Bourguignon et al., 2020) and either an increase or a decrease in effective population size *N*_e_. While some observations link the decrease in genome size to the increase in random drift (Moran, 2002; Nilsson et al., 2005; Kuo et al., 2009), this is not consensual among the scientific community since long term reduction in *N*_e_ is also thought to increase genome complexity and genome size: the increase in genetic drift would cause the fixation of slightly deleterious duplications, which would be more frequent than slightly deleterious deletions (Lynch and Conery, 2003; Lefebure et al., 2017). Deletion biases, or more generally the balance between insertion and deletion rates and spectra, may also play a role in genome size evolution (Petrov, 2002). In particular, deletion biases are believed to contribute to the small genome size of prokaryotes (Bingham and Ratcliff, 2024). However, the interaction of various mutational biases with changes in mutation rates, or with changes in effective population size, has yet to be investigated in more details.

In this study, we focus on determining the impact of both an increased mutation rate and a change in population size on genome size evolution. However, mutation rates and population sizes are difficult to estimate. The effective population size is also highly variable throughout time, such that it is not totally obvious which long-term average is relevant at the macroevolutionary scale (Brevet and Lartillot, 2021; Müller et al., 2022). For that reason, many comparative analyses have relied on somewhat indirect proxies, such as life-history traits (Popadin et al., 2007; Romiguier et al., 2012; Figuet et al., 2016). However, the precise quantitative relation between these proxies and effective population size is unknown. Moreover, the very different living conditions and potential mutational biases of the bacterial species that have undergone genome reduction introduce many confounding factors. To avoid these pitfalls, we choose to turn to simulation to study the different phenomena leading to genome size reduction, and more specifically the influence of population size and mutation rate over the coding and non-coding genome sizes, with or without mutational biases.

*In silico* experimental evolution provides tools to study genomic architecture in detail (Adami, 2006; Hindré et al., 2012; Batut et al., 2013). For our study, we need a framework that provides coding and non-coding genomic compartments which can vary independently, and with arbitrary underlying mutational biases for the deletion/insertion balance. Then, running simulations in a perfectly controlled environment covering a broad range of populations sizes *N* and mutation rates *µ* makes it possible to investigate the conditions and mechanisms leading to genome size reduction. We will hence use Aevol, a simulation platform that provides an explicit genomic structure where both the coding and non-coding genome can evolve freely. Aevol emulates the evolution of bacteria and enables replicated and controlled *in silico* evolution experiments with known and fixed parameters (Knibbe et al., 2007; Banse et al., 2023). It provides an ideal tool to uncover links between genome size and either population size or mutation rate, as the experimenter perfectly controls these parameters. Throughout the experiments, fitness, genome size and amounts of coding and non-coding bases are monitored to study the evolution of genome architecture and the response of genome size to changes in *µ* and *N*.

Our results show that both an increase in *N* or *µ* lead to genome size reduction, regardless of the underlying mutational bias. However, both conditions lead to very different genome structures, as a high *µ* reduces both coding and non-coding compartments while a high *N* reduces only the non-coding compartment, resulting in densely packed genomes. Surprisingly, they nevertheless both lead to a similar coding proportion when increased by the same factor, such that *N × µ* appears as a key compound parameter determining genome structure. Finally, by measuring the robustness of the lineages evolved in different conditions, we show that the observed variation in genome size and structure are due to the interaction between selection for phenotypical adaptation to the environment and selection for robustness.

## 2 Results

We perform our experiments using Aevol, a forward-in-time evolutionary simulator (Knibbe et al., 2007; Banse et al., 2023). Aevol is an individual based model which includes an explicit population and in which every organism owns a double-stranded genome. It uses an explicit genome decoding algorithm directly inspired from the central dogma of molecular biology, to compute the phenotype, and thus the fitness, of each individual based on its genomic sequence. As Aevol also includes a large variety of mutational operators (including substitutions, InDels and chromosomal rearrangements), this non-parametric genotype-to-phenotype map allows for changes in the genome architecture (genome size, coding density, overlapping genes or operons, etc.), without assuming a predefined distribution of fitness effects. Indeed, in the model, it is possible to reach similar fitnesses in many ways, by adjusting the number of genes, their loci, their lengths, or the intergenic distances, hence the total amount of non-coding DNA. In Aevol, genes are typically created by duplication-divergence (Kalhor et al., 2024), but they can also be deleted, and some emerge *de novo*. Hence, the impact of a given mutation highly depends on the preexisting genome structure, which can in turn be indirectly selected (Knibbe et al., 2007). Aevol therefore allows studying changes in size and structure of genomes in response to changes in population size and mutation rates.

Our experiments start from five “Wild-Type” (WT) lines, each having evolved for 10 million generations within a population of 1, 024 individuals and a mutation rate of 10^*−*6^ mutations per base pair for each mutation type: substitutions, small insertions, small deletions, duplications, deletions, translocations, and inversions. There is no underlying mutational bias: the insertion and deletion of bases are equally probable. The five WTs display stable genome structures (with small random variations, as exemplified by cases *N*_0_ and *µ*_0_ on Figs. 1 and 2) although they still slowly gain fitness by fixing rare favorable mutations (see case *N*_0_ on Figure 5A). Their fitness and genomic characteristics are displayed in section 4.2, Table 1. In our experiments, these WTs are used as founders of new populations, which are confronted to new evolutionary conditions for 2 million generations. In parallel, these same WTs were evolved in the same conditions the WTs were evolved in, providing perfect control experiments. We compare the fitness, genome size and genome structure of populations evolved in new conditions with those of the control populations. Finally, we repeat part of these experiments with WTs that evolved with either an insertion or a deletion bias to understand how an underlying mutational bias might impact our findings.

**Table 1:** Characteristics of the 5 Wild-Types at the start of our experiments.

| WT id | Fitness<br>(arbitrary unit) | Total genome size<br>(bp) | Coding size<br>(bp) | Non-coding size<br>(bp) | Coding<br>fraction |
| --- | --- | --- | --- | --- | --- |
| 1 | 0.014903 | 13,599 | 9,395 | 4,204 | 0.69 |
| 2 | 0.103795 | 13,660 | 8,828 | 4,832 | 0.65 |
| 3 | 0.128472 | 14,171 | 9,507 | 4,664 | 0.67 |
| 4 | 0.035369 | 14,507 | 10,003 | 4,504 | 0.69 |
| 5 | 0.029588 | 14,290 | 10,644 | 3,646 | 0.74 |
| Average | 0.0624254 | 14,045.5 | 9,675.4 | 4,370 | 0.69 |

**Figure 1.**
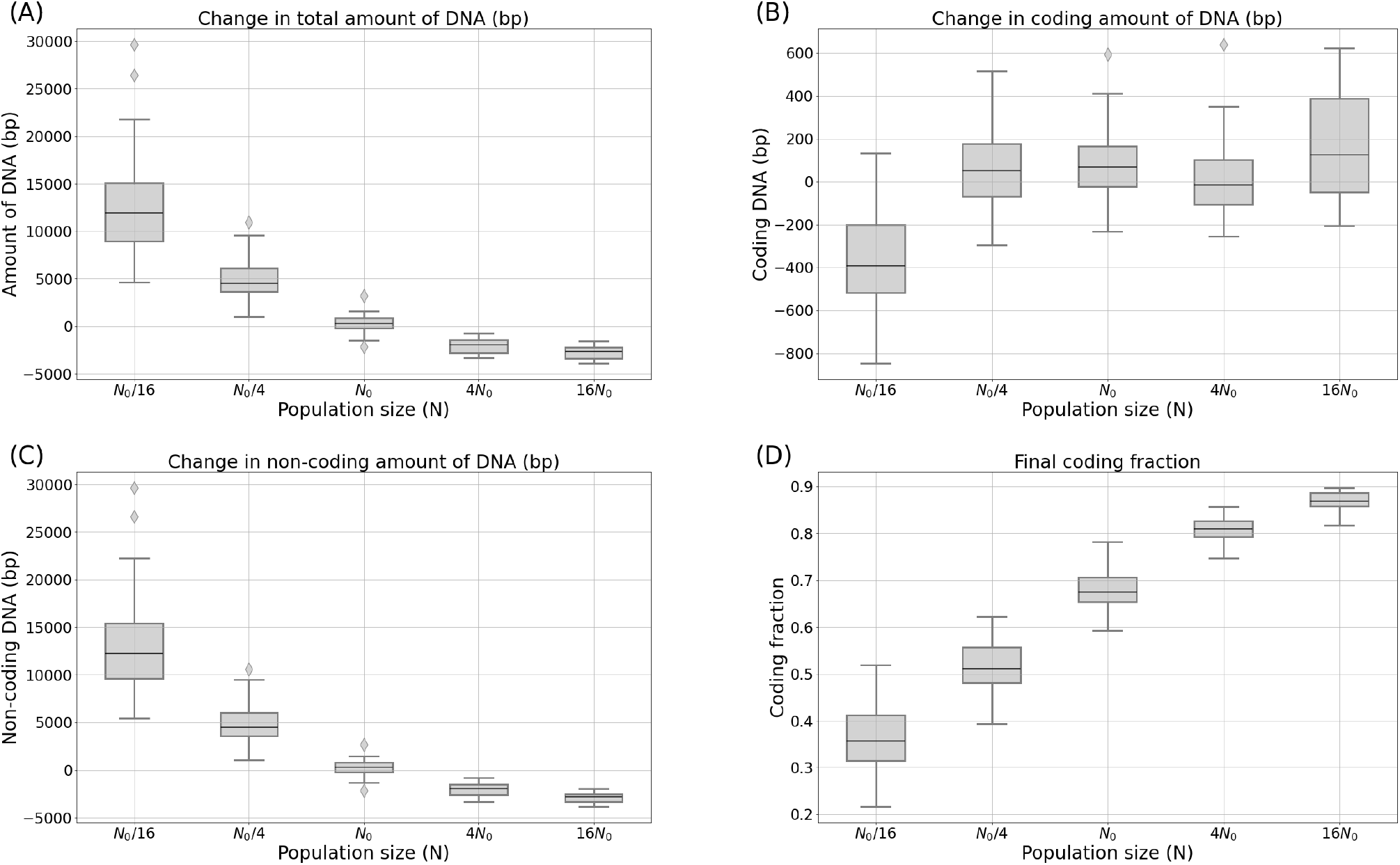
**Total (A), coding (B) and non-coding (C) genome size variation, and final coding fraction (D)**, after 2 millions generations. For each of the 5 WTs, 10 replicas were performed under a constant mutation rate (*µ*_0_ = 10^*−*6^ per base pair for each type of mutation) with 5 different population sizes (*N*_0_ = 1, 024 being the control population size).

**Figure 2.**
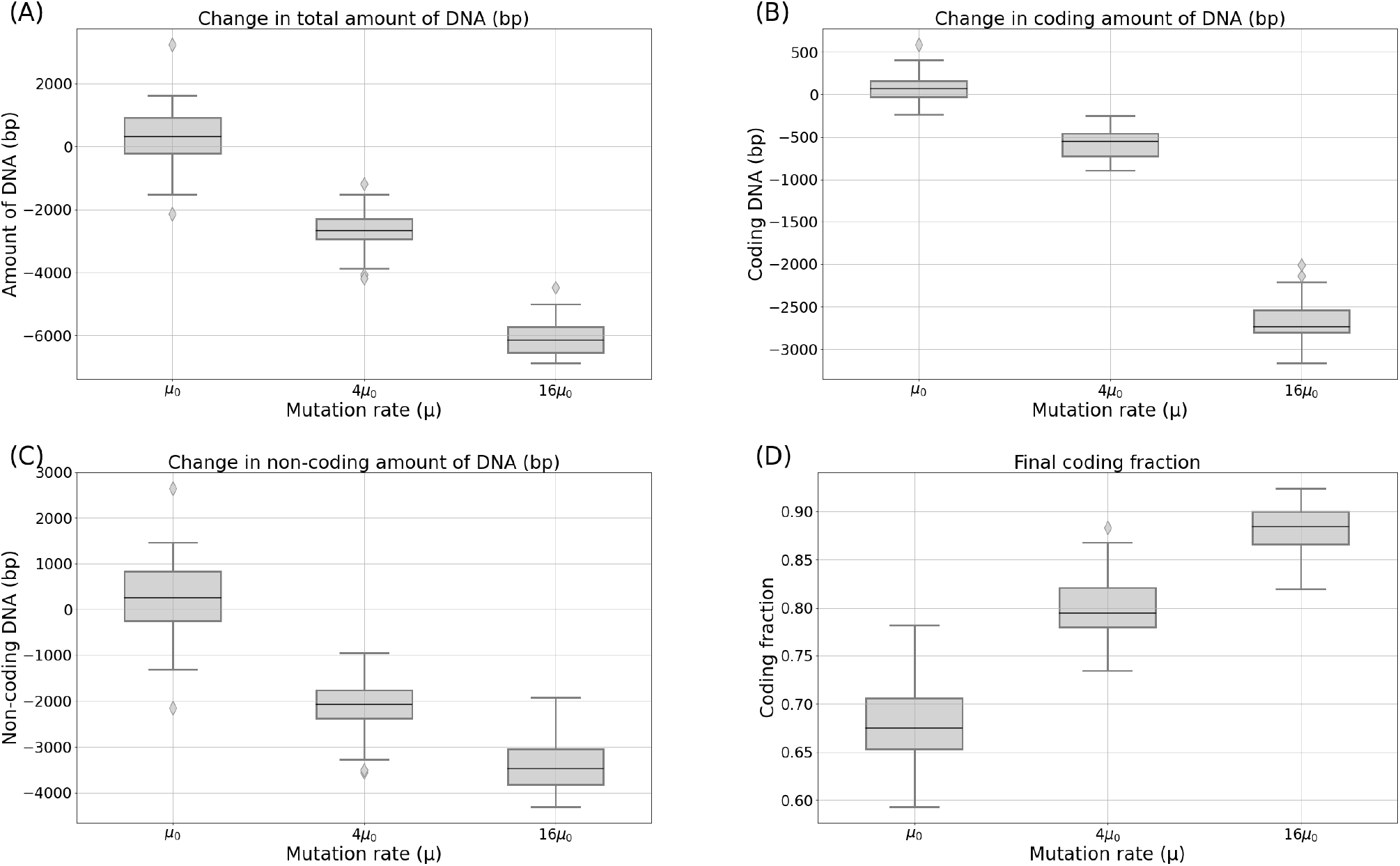
**Total (A), coding (B) and non-coding (C) genome size variation, and final coding fraction (D)**, after 2 millions generations. For each of the 5 WTs, 10 replicas were performed under a constant population size (*N*_0_ = 1, 024 individuals) with 3 different mutation rates: the control *µ*_0_ = 10^*−*6^ mutations per base pair for each type of mutation, 4 *× µ*_0_ and 16 *× µ*_0_.

### 2.1 Genome size evolution following a change in population size and mutation rate

#### 2.1.1 Change in population size

In the absence of mutational bias, increasing the population size by a factor 4 or 16 results in a reduction in the total genome size (see Figure 1A). Yet, this change does not impact the coding and non-coding parts of the genome proportionally: while the size of the coding compartment is barely affected (see Figure 1B), the non-coding genome size is greatly reduced (see Figure 1C). As a result, the coding proportion of the genome increases (see Figure 1D). Conversely, reducing the population size by a factor 4 or 16 increases the total genome size (Figure 1A) by increasing greatly the non-coding genome size (Figure 1C). In the extreme condition *N*_0_*/*16, the coding genome size is also slightly reduced (Figure 1B). As a result, the coding fraction of the genome is drastically reduced (Figure 1D).

#### 2.1.2 Change in mutation rate

In the absence of mutational bias, increasing the mutation rate drastically reduces the total genome size (see Figure 2A). Thus, at first sight, population size and mutation rate seem to have a similar effect on genome evolution. However, in the details, the effect of these two variables on genome structure appears to differ, as the reduction now occurs in both the coding and noncoding genomic compartments (see Figure2B and C). Both are nevertheless not proportionally affected by the decrease in mutation rate which affects more strongly the non-coding part of the genome, such that the final coding fraction of the genome increases with *µ* (see Figure 2D). Altogether, these results show that streamlined genomes, denser and shorter than their ancestors, can result from either an increase in population size or in mutation rate.

Notably, and in spite of the very different dynamics displayed in the two experiments, a 4-fold increase in *N* or in *µ* results in the same final coding proportion of approximately 80%. The same is true for a 16-fold increase (88%). To further investigate this result, we conducted additional experiments to observe the combined effects of a simultaneous modification in both *N* and *µ*.

#### 2.1.3 Linked effect of population sizes and mutation rates

Figure 3 shows the variation in the total amount of DNA, coding size and non-coding size, as well as the variation in coding fraction for several combinations of changes in *N* and *µ* (note that, in the panels of Fig 3, the bottom line and the central column respectively correspond to the values presented in Figs. 1 and 2).

**Figure 3.**
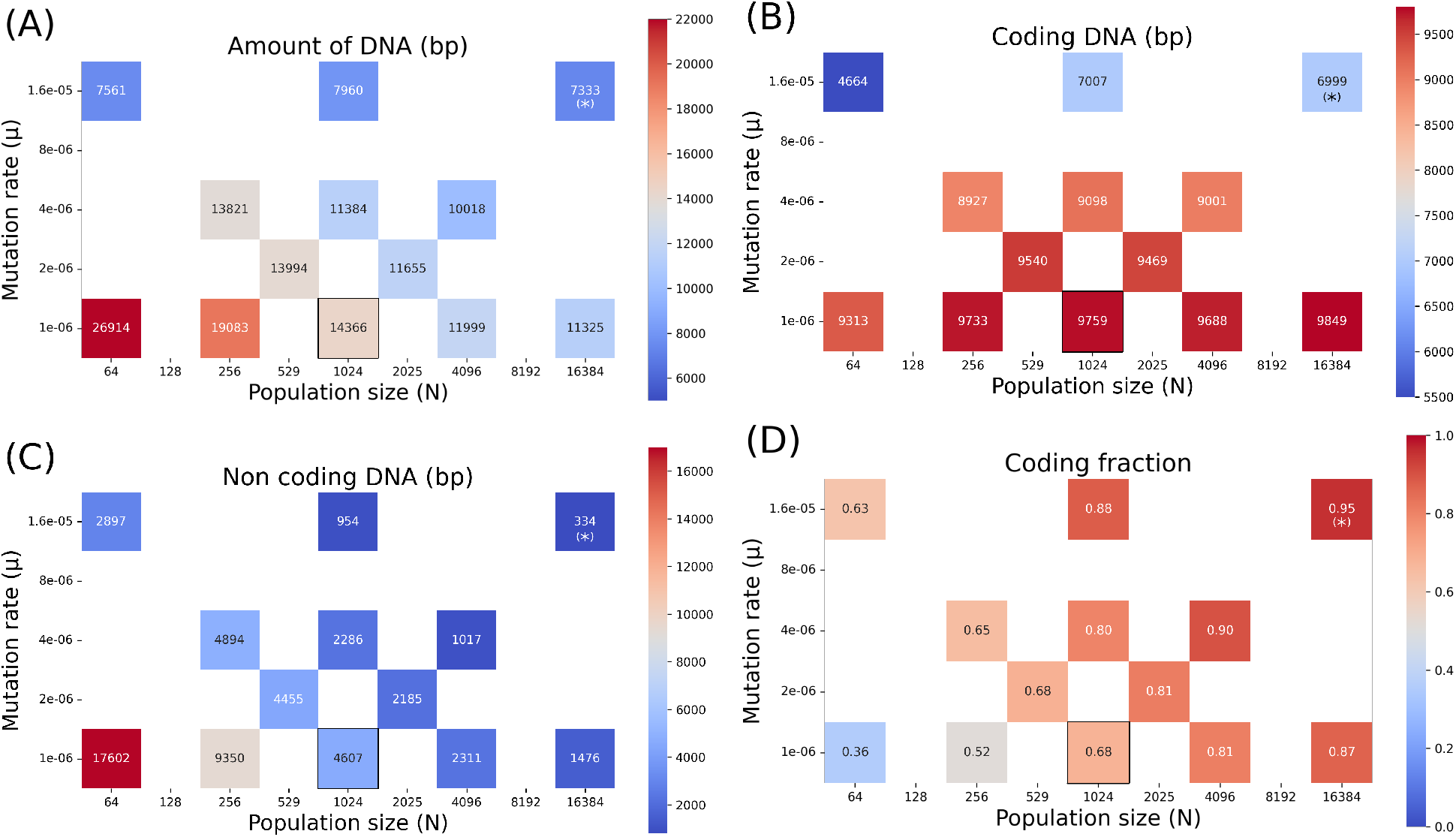
**Amount of DNA (A), coding size (B), non-coding size (C) and coding fraction (D)** for the different combinations of *µ* and *N* tested, after 2 million generations. For each of the 5 WTs, 10 replicas were performed for each tested set of conditions. Control conditions (*N* = 1, 024 and *µ* = 1.10^*−*6^) are outlined in black. For the combination of both the highest mutation rate and the largest population size, only the median was tested due to computational limitations, which is indicated by a (*).

Overall, as *N* increases, the total amount of DNA decreases, whatever the value of *µ* (see Figure 3A). A higher *µ* also leads to a reduction in the total genome size, whatever the value of *N*. However, the effect of population size and mutation rate differ when considering the coding size of the genome: specifically, the coding size increases with *N* but decreases with *µ* (see Figure 3B). This is counterbalanced by the change in the non-coding size of the genomes (see Figure 3C), which strongly decreases with both *N* and *µ* and drives the overall change in genome size.

The interplay between *N* and *µ* results in a surprisingly constant coding fraction across the different constant values of *N × µ* (see Figure 3D). Indeed, we observe that under constant *N × µ*, and despite the fact that these two factors taken individually have changed in different proportions, the coding fraction remains constant: 80% when *N*_0_ *× µ*_0_ is multiplied by 4 compared to the control conditions, and 88% when *N*_0_ *× µ*_0_ is multiplied by 16 (see Figure 3D). Although the coding fraction does slightly vary (from 68% to 63%) for the most extreme change tested (*N*_0_*/*16 and 16*µ*_0_), the diagonals of constant *N*_0_ *× µ*_0_ also display an almost constant coding fraction (Figure 3D).

However, strikingly, the total genome size as well as the coding and non-coding genome sizes vary greatly, even for similar coding densities (Figure 3B, C and D). For densities of 63% and 65%, the total amount of DNA can be almost halved (from 13, 821 bp to 7, 561 bp) by going from *N*_0_*/*4 and 4*µ*_0_ to *N*_0_*/*16 and 16*µ*_0_ on the same diagonal of constant *N × µ*. Conversely, we can reach similar values of genome size (11, 300 bp) despite important differences in the coding percentage (80% when *µ* is multiplied by 4, and 87% when *N* is multiplied by 16). Altogether, these results show that a large range of genome sizes and structures (here corresponding to coding densities) can result from a combined variation in both the population size *N* and the mutation rate *µ*.

### 2.2 Mutational biases change the equilibrium genome size, but not the role of *N* and *µ*

As genome sizes are generally thought to be heavily impacted by mutational biases, we control whether the effect of population size and mutation rate we observed is affected by either a deletion or an insertion bias. To this end, we evolved 5 Wild-Type organisms with either an insertion bias (twice as many duplications than large deletions), or a deletion bias (twice as many large deletions than duplications). The rates of all other types of mutations, as well as the sum of all mutation rates, are the same as in the previous experiments. As expected, the equilibrium genome sizes and coding proportions of these Wild-Types is affected by the balance between large deletions and duplications, with an average genome size of 11, 623 bp in the presence of a deletion bias, and 16, 350 in the presence of a duplication bias (instead of 14, 046 bp without any bias). The coding proportion is also affected: 0.78 and 0.61 respectively, instead of 0.69. This shows that the genome size and structure are, as expected, strongly influenced by the underlying mutations biases.

We then confronted the median (in terms of genome size) WT of each condition to changes in population size (multiplied or divided by 4) or mutation rate (multiplied by 4) for 10 replicas. Similarly to what is observed without bias, an increase in *N* reduces the non-coding genome size only, while an increase in *µ* reduces both the coding and non-coding genome (see Figure 4). Notably, a decrease in *N* increases the non-coding genome size, even in case of a deletion bias, although an insertion bias greatly amplifies this effect. As a result, and despite the strong mutational biases, we observe that multiplying either the population size or the mutation rate by the same factor lead to a genome compaction in similar proportions (the final coding fraction being 0.85 vs. 0.88 in case of the deletion bias, and 0.78 vs. 0.77 in case of the insertion bias respectively). Therefore, although mutational biases influence the equilibrium genome sizes and structures, they do not fundamentally change how the genomes react to variations in population size or mutation rate. In other words, our simulations show that mutational biases only determine the equilibrium set point around which population size and the overall mutation rate then modulate the genome size and structure. Similar experiments were run with biases in InDels and are presented in the supplementary materials S2.

**Figure 4.**
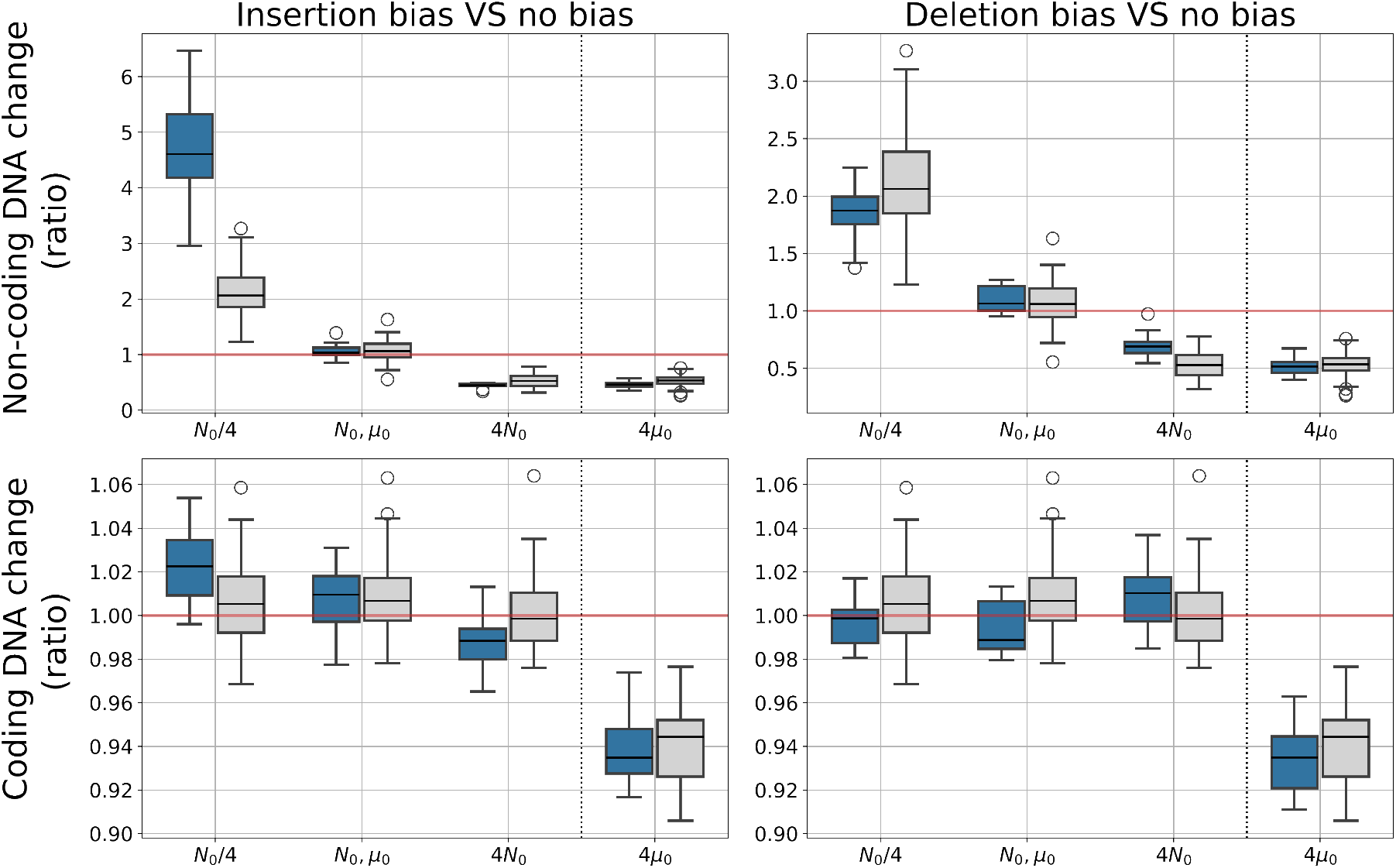
Change in coding and non-coding genome sizes in reaction to changes in *N* or *µ* for the different mutational biases. Blue boxes show the results with a mutational bias (left: insertion bias, right: deletion bias), and gray boxes show the results without mutational bias. Depicted values are the ratio of the coding/non-coding sizes at the final generation over the value at generation 0.

### 2.3 Robustness selection as the explanatory mechanism

We observed that two distinct processes, triggered by an increase in either population size or mutation rate, can lead to genome size reduction in our experiments. However, both have different effects on coding and non-coding sequences: while an increased *µ* reduces both the coding and non-coding genome sizes, increasing *N* reduces only the non-coding genome size.

We propose that these observations can be explained by an interplay between selection for phenotypic adaptation to the environment (hereafter *direct selection*), and selection for replicative robustness (hereafter *indirect selection*). We define the replicative robustness of an individual as the proportion of its offspring that did not acquire new deleterious mutations. In our case, wild-type organisms are very well adapted to their environment, thus most mutations will be strongly deleterious if they affect the coding part of the genome. This is particularly true for chromosomal rearrangements that can affect large genomic segments. Conversely, beneficial mutations are extremely rare. We therefore approximate the robustness of our organisms by measuring the proportion of their offspring that have the exact same fitness, *i.e*. that underwent no mutations or only neutral mutations.

A more robust individual has more chances to pass on its genomic information accurately than a less robust one, thus enabling its lineage to better maintain its fitness in the long term, and to outcompete other lineages in which deleterious mutations would accumulate at a higher rate. This results in an indirect selection for replicative robustness. Importantly, replicative robustness depends both on the probability for a given mutation to be neutral (hence on the fraction of non-coding sequences in the genome) and on the mean number of mutations underwent by the genome at each generation (hence on the total amount of DNA). By contrast, direct selection depends only on the content of the coding compartment, the size of which is likely to be positively correlated with the level of phenotypical adaptation (at least in our model). As a result, indirect selection for robustness favors shorter genomes with a lower coding fraction, while direct selection for phenotypical adaptation maintains or even increases the coding size of the genome.

The efficacy of both direct and indirect selection increases with population size, since some deleterious mutations that were quasi-neutral for a low *N* can become effectively counterselected in the context of a high *N*, changing the balance of beneficial *vs* deleterious fixed mutations. To quantify this effect, we measured the robustness of the individuals at time 2, 000, 000 in the simulations without mutational biases. Figure 5A and 5B show that the increase in selection efficacy indeed induces both an increase in fitness (due to direct selection) and an increase in replicative robustness (due to indirect selection). In terms of genomic structure, a more efficient direct selection (*i.e*. a weaker random drift) is thus expected to increase the coding genome size, and a more efficient indirect selection is expected to decrease the overall genome size. The combination of both these effects leads to a decrease in the noncoding genome size, and a maintenance of the coding genome size as exemplified by Figure 1B and C. Conversely, when the population size is reduced, the increased drift leads to the loss of coding sequences and an inflation of the non-coding compartment (Figure 1B and C). This reorganization of the genome structure is associated with a loss in robustness (Figure 5B).

**Figure 5.**
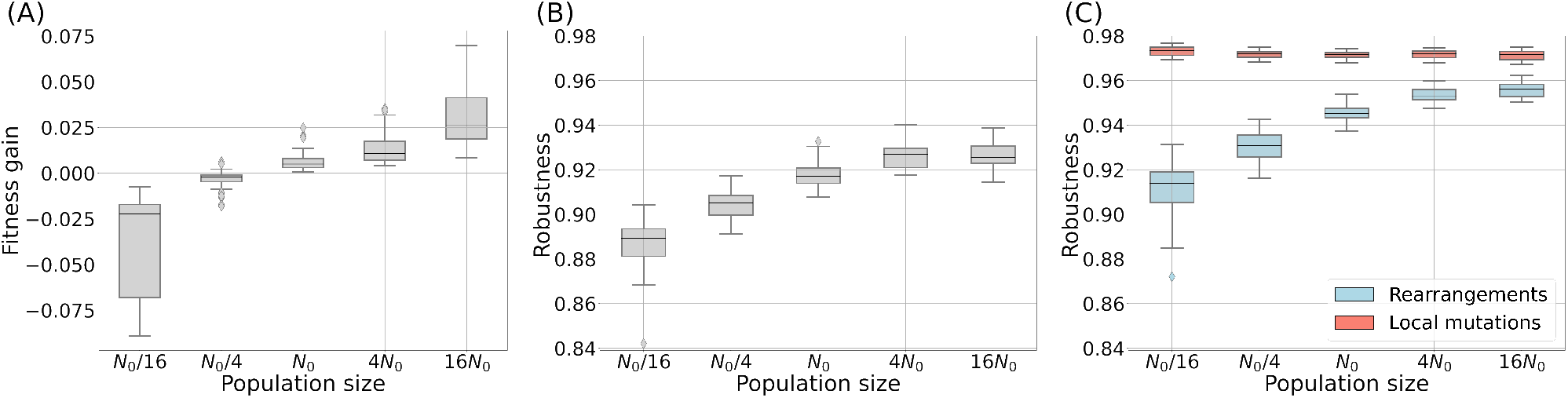
Fitness gain (A) and Robustness (B: overall and C: by mutation type) at the end of the simulations, for different population sizes *N* and without mutational biases. Robustness is defined as the proportion of neutral offspring. The mutation rate is fixed to 10^*−*6^ per base pair for each type of mutation.

In Aevol, genomes undergo different types of mutations that can be roughly grouped into local mutations (substitutions, InDels) and chromosomal rearrangements (duplications, deletions, inversions, translocations). Both kinds of events don’t have the same effect on robustness. Figure 5C shows the change in robustness induced by the different types of events. It shows that the loss and gain in robustness are driven by chromosomal rearrangements. In contrast, local mutations (substitutions and InDels) do not have a significant effect on robustness.

In case of an increased mutation rate, things are very different: a sudden increase in *µ* results in an immediate drop in robustness at the beginning of the experiments (Figure 6A). As the proportion of offspring that bears mutations rises with *µ*, we go from an initial robustness of 92% for *µ*_0_, to 71% for 4*µ*_0_, and only 26% for 16*µ*_0_. In these new conditions, organisms are no longer able to transmit their genome to the next generation without deleterious mutations, and thus the indirect selection for robustness becomes temporarily stronger than the direct selection for phenotypical adaptation. Indeed, features that would not be accurately inherited cannot be selected. This indirect selection for robustness leads to the fixation of mutations that drastically decrease the genome size, even at the cost of a loss of fitness for the individuals (see Figure 6B): the only lineages that survive in the long term are those that have undergone a decrease in genome size, allowing them to mitigate the effect of the per-base mutation rate, thus regaining some robustness (see Figure 6C). Once the robustness has increased sufficiently, direct selection for phenotypical adaptation can resume and the fitness starts to increase again (see Figure 6B). Interestingly, organisms manage here to continue to lose some coding base pairs while increasing their fitness, probably thanks to global genome restructurings allowing for a more compact encoding of the phenotype, for example through overlapping genes. This dynamic is very different to what happens when *N* is increased (and so the initial robustness is unaffected), as shown by Figure 6D, E and F.

**Figure 6.**
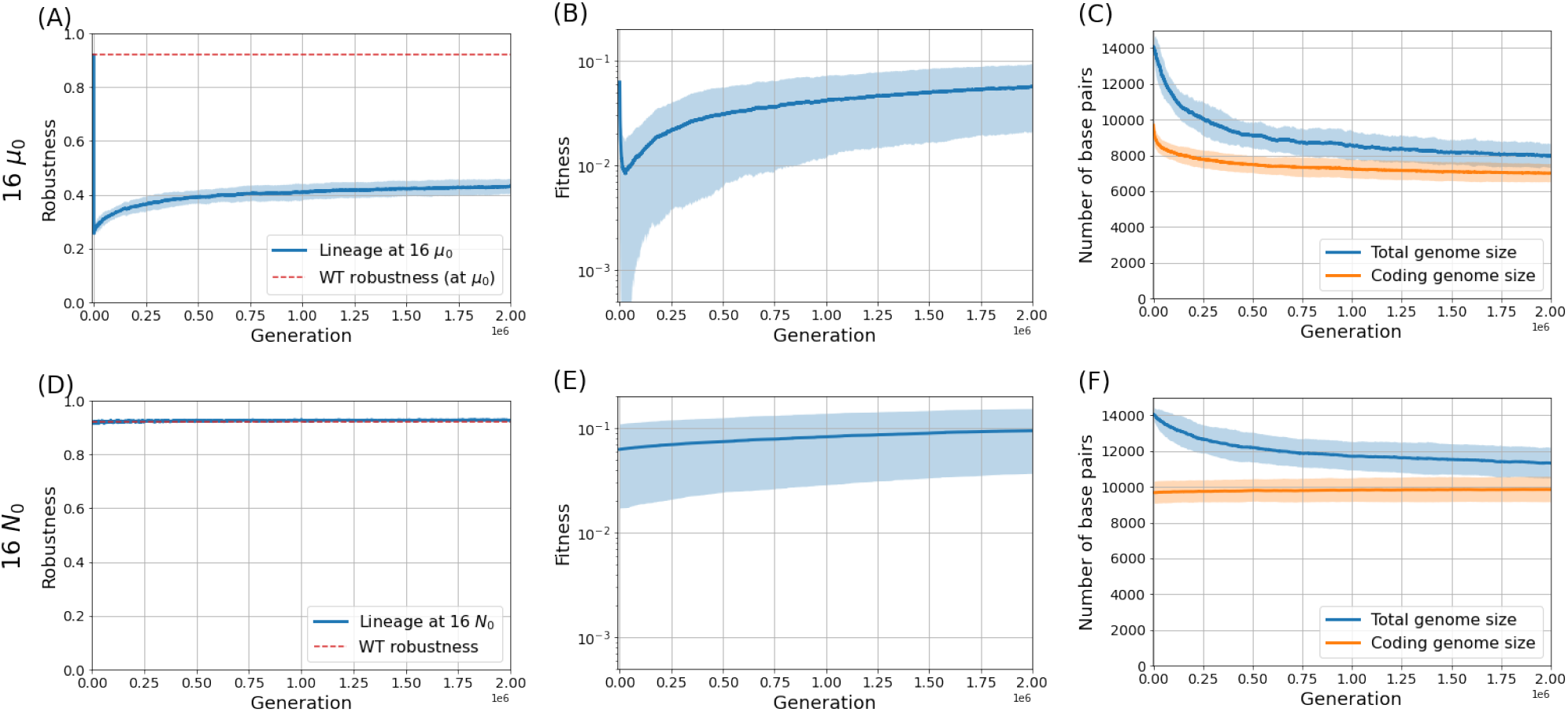
**Robustness, fitness, and genome architecture** across generations for *µ* = 1.6 *×* 10^*−*5^ (16 *µ*_0_) per base pair for each mutation type and *N* = 1, 024 (*N*_0_) (top row, panels A, B and C) and *N* = 16, 384 (16 *N*_0_) and *µ* = 1 *×* 10^*−*6^ (*µ*_0_) per base pair for each mutation type (bottom row, panels D, E and F). Lines represent the mean values across the 50 simulations, and the shaded areas represent the standard deviations.

Notably, robustness does not reach values as high as before the increase in mutation rate and stays below 50%. Indeed, the genome size could not be divided by 16 while keeping a good enough phenotypical adaptation, and the selection for phenotypical adaptation becomes stronger than selection for robustness as soon as some organisms are able to pass on their genomic information.

The interplay between direct and indirect selection can therefore explain both types of genome size reduction: affecting both coding and non-coding compartments (although not proportionally) when caused by an increased mutation rate, and focused on the non-coding compartment when caused by an increased population size.

## 3 Discussion

We found that, in our experiments, genome size reduction can be caused by an increase in population size, in mutation rate, or both, even in case of mutational biases. These two factors can nevertheless be distinguished, as they have different effects on the coding and non-coding sequences of the genome. Their combination in various proportions can create a broad range of alternative patterns of genome size and coding density. In particular, by playing independently on mutation rate and population size, our model can reproduce the two extreme but different cases of genome size reduction that are seen in some endosymbionts and cyanobacteria. As an example, *Prochlorococcus marinus* is known to have lost both some parts of its coding and non-coding genome, although in different proportion such that its coding density has increased (Dufresne et al., 2005; Batut et al., 2014; Giovannoni et al., 2014). In our model, this would correspond to a population undergoing an increase in population size and a slight increase in mutation rate, which is coherent with the scientific literature on *Prochlorococcus marinus* (Hu and Blanchard, 2008; Marais et al., 2008), although the effective population size of this species has been recently questioned (Chen et al., 2022). On the other hand, *Buchnera aphidicola* has conserved its coding proportion but greatly reduced its total genome size (Moran and Mira, 2001), which could be explained in our model by an increase in mutation rate and a decrease in population size, in similar proportions. This suggests that indirect selection for shorter genomes through robustness selection could be a key factor of genome evolution (Wilke et al., 2001; Gabzi et al., 2022), and especially of the evolution of genome size and structure.

Our observations confirm those made by Lynch and Conery (2003) that an increased genetic drift, here associated with a decreased population size, increases the genome size. Our result also point towards an equilibrium genome size: a sufficient number of genes makes it possible to fine-tune the phenotype to the environment, but the genome also has to be short enough to prevent the degeneration caused by an excess of chromosomal rearrangements (Knibbe et al., 2007; LaBar and Adami, 2020). Increasing the mutation rate or the population size displaces this equilibrium toward shorter genomes, either through a more efficient genome purification of non-coding sequences (when increasing *N*) or a loss of both coding and non-coding sequences to recover a minimal level of robustness (when increasing *µ*). Of course, mutational biases (in terms of the balance between insertions and duplications versus deletions) also play an important role in determining the equilibrium genome size. In particular, deletion biases have been suggested as one main reason explaining why bacterial genomes remain small (Mira et al., 2001). However, what we show here is that, because of indirect selection for robustness, a deletion bias is in fact not needed to prevent a runaway inflation in the size of genomes. Instead, selection for robustness provides a counteracting force – which increases with genome size, so it should eventually offset an underlying bias in favor of insertions or duplications. Importantly, this indirect selection was not postulated in the model but emerged spontaneously in the simulations. We propose an evolutionary mechanism consisting of a trade-off between direct selection for phenotypical adaptation and indirect selection for replicative robustness. In that aspect, mutations appear to be a weak selective force, as pointed by Lynch and Walsh (2007). However, the emphasis was previously on the mutational targets contributed by genomic features, such as introns. Here, we emphasize another aspect, which seems to have been overseen thus far: any non-functional DNA represents an additional target for initiating macroscopic mutational events. This mechanism requires no additional hypotheses and is very general. It should therefore be pervasive in the living world. We do nevertheless expect some other determinants of genome size, such as insertion sequences or effects of the non-coding genome over gene expression, to have an effect on genome size variation. These could be added to the model in order to study their impact on top of the mechanisms presented here. However, we argue that they would not fundamentally compromise the dynamics observed here and only displace the final equilibrium.

Although our main focus was on the final equilibrium reached by the populations after a change in *N* or *µ*, our observations are broader than the end equilibrium, as we can observe the temporal dynamics (Figure 6 and S3 to S15). In particular, we observe that, when the mutation rate increases strongly, the fitness initially drops drastically (Figure 6B). This can be related to an error-threshold crossing mechanism (Eigen, 1971; Takeuchi and Hogeweg, 2007; de Boer and Hogeweg, 2010): individuals can no longer pass on to their descendants all the information contained in their genome. They therefore lose fitness, and the lineage that survives in the long term is the one where genomes greatly reduced in size in the early phase of the experiment, thus reducing the number of mutations per replication event and finally reaching a point at which the information can be passed on reliably. The detailed aspects of these temporal dynamics could be the focus of future work. Indeed, it has been shown that genome reduction in endosymbionts occurred very quickly after the endosymbiosis became effective (Moran, 2003; Wernegreen, 2015), which is also what we observed in our data (Figure 6).

In our experiments, *N × µ* stands out as a determining factor of genome structure, as isoclines of identical *N × µ* values display similar coding densities, even in case of reduced genomes or mutational biases. Understanding this invariant is one of the most exciting perspectives opened by our work. Its importance had already been highlighted by Lynch et al. (2006) in organelles, but our results suggest that this joined factor of drift and mutational pressure is determinant to genome evolution throughout the tree of life. Notably, there is a small variation in coding fraction along our *N* isoclines, which could be due to our use here of population size (*N*) instead of effective population size (*N*_e_). Indeed, in our setup, the competition is local and thus *N*_e_ is slightly greater than *N*, but this relationship is not linear (see Supplementary Data S1). Further versions of the model could rely on various measures of the effective population size to reach more accurate predictions, but we believe that our results can be interpreted nonetheless, as changes in population size and in effective population size are very similar over the range of population sizes tested here (see Supplementary Data S1).

Naturally, various other mechanisms can have an impact on genome size evolution. For instance, there can be a limitation in available resources for nucleotides production, constraining the total genome size (Ngugi et al., 2023). Transposable Elements (TE) have also been shown to induce genome streamlining because genome reduction prevents TE invasion (van Dijk et al., 2022). Bacterial recombination could also further complicate the picture by adding a new type of mutation with unexpected interactions. It would be very interesting to add all of these additional features to the model in order to study their impact on genome size evolution. They would contribute to new mechanisms and pressures on genome size and structure, changing the equilibrium set point. However, our results suggest that the qualitative outcome of the balance between phenotypic selection and indirect selection for robustness that we highlight here would be robust to addition of these mechanisms.

Our study also raises the question of the generalization of the relationships observed here between *N, µ* and the genome structure to eukaryotes. Indeed, Aevol is fundamentally a model of the evolution of prokaryotic genomes. The introduction of sexual reproduction and meiotic recombination to our model could have unexpected consequences, and may thus lead to qualitatively different results. The robustness of our observations to such big changes is yet to be assessed, and it would be interesting to study the interplay between selection for phenotypical adaptation and selection for robustness in the presence of sex and meiotic recombination.

To conclude, our experiments show that genome size reduction can occur in two very different conditions for bacteria. On the one hand, a very large population size promotes a more efficient selection in face of random drift, which in turn enhances the robustness of genomes by decreasing their non-coding load. This corresponds to streamlining, and leads to genomes with a high coding density. On the other hand, a higher mutation rate results in an instantaneous decrease in the robustness of genomes in the entire population, making the selection for robustness transiently stronger than the selection for phenotypical adaptation. The genome then shrinks rapidly, with both coding and non-coding sequences being discarded until a new robustness equilibrium is reached, all this at a substantial initial cost in phenotypical adaptation. This corresponds to a decaying genome and is compatible with empirical observations in endosymbiotic bacteria (Moran, 2003). Strikingly, this remains true even in the presence of a mutational bias. Although the model that we propose here, of a balance between selection for robustness and selection for phenotypical adaption, can explain the tendencies we observe and the final genome structures in our populations, further work is needed to understand the transient regimes and the mechanisms behind the constant coding fraction along the *N × µ* isoclines.

## 4 Materials and Methods

### 4.1 The Aevol framework

Aevol (Knibbe et al., 2007; Banse et al., 2023) is an individual-based forward-in-time simulation software that has been specifically designed to study the evolution of genome structure. It emulates a population which is composed of a fixed number of individuals on a grid (Figure 7A). Each individual owns a double-stranded circular genomic sequence, composed of 0s and 1s. In order to compute the phenotype, sequences on the genome are recognized as promoters and mark the start of transcription, which stops when a sequence able to form a hairpin structure is encountered. On RNAs, Shine-Dalgarno-like sequences followed by a START codon mark the beginning of translation. The RNA sequence is then read 3 bases at a time until a STOP codon is encountered on the same reading frame. An artificial genetic code allows for each sequence of codons to be converted into a mathematical function, and the sum of all functions encoded on the genome defines the phenotype of the individual (Figure 7B). The distance between this function and a target function, which represents the ideal phenotype in the specified environment, gives the fitness of the individual with a scaling factor *k* that tunes the strength of the selection. A detailed explanation can be found on the dedicated website www.aevol.fr.

**Figure 7.**
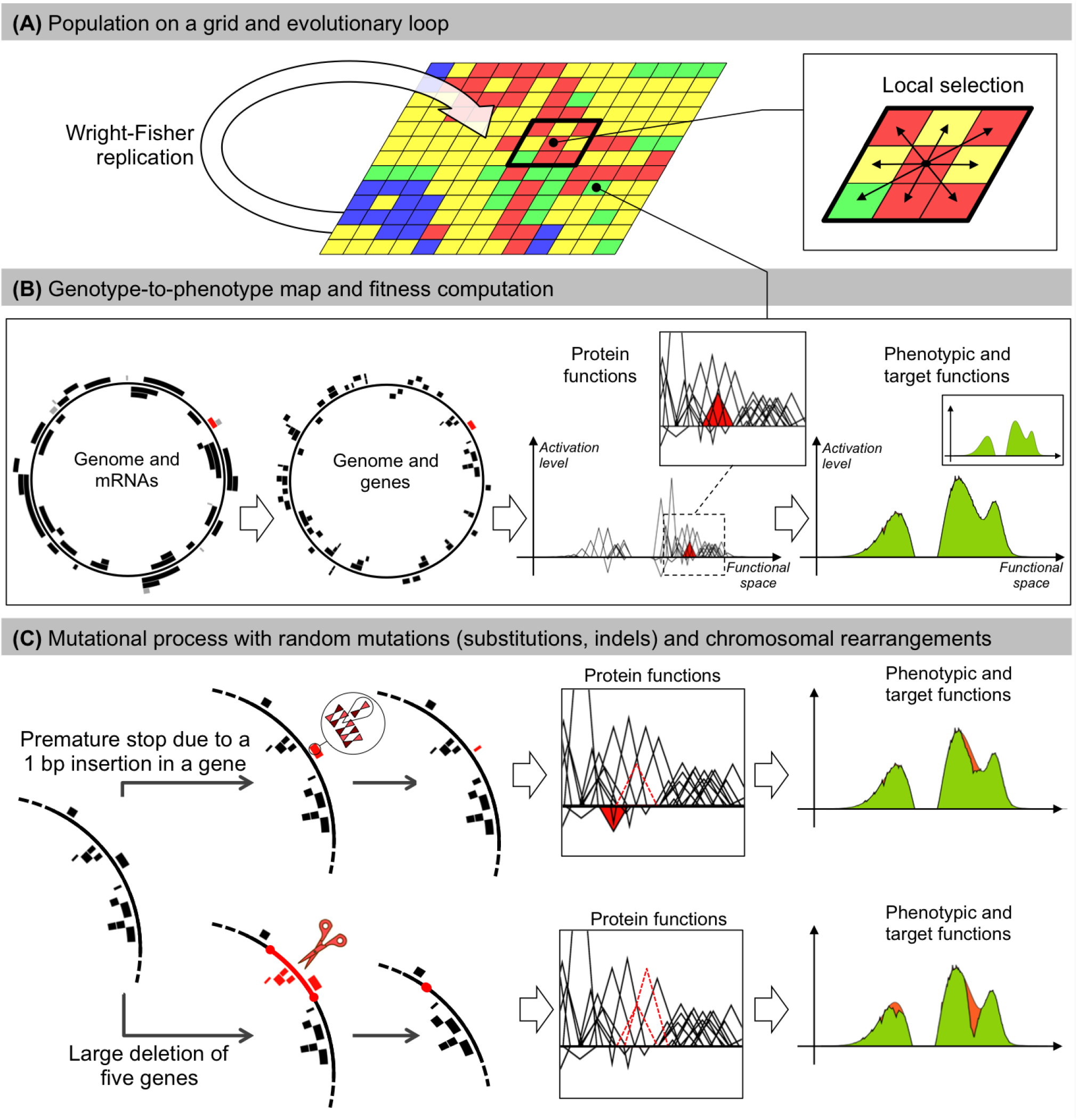
The Aevol model. **(A)** Individuals are distributed on a grid. At each generation, the whole population replicates according to a Wright-Fisher replication model, in which selection operates locally within a 3*×*3 neighborhood. **(B)** Each grid-cell contains a single organism described by its genome. Genomes are decoded through a genotype-to-phenotype map with four main steps (transcription, translation, computation of protein functions, computation of the phenotype). Here, for illustration purposes, a random gene and the corresponding mRNA are colored in red. The red triangle represents the function of this gene in the mathematical world of the model. The phenotypic function is calculated by summing all protein functions. The phenotype is then compared to a predefined target (in green) to compute the fitness. The individual presented here has evolved in the model during 500, 000 generations. **(C)** Individuals may undergo mutations during replication. Two example mutations are shown: A small insertion (top) and a large deletion (bottom). Top: A 1 bp insertion occurs within a gene. It causes a frameshift, creating a premature stop codon. The ancestral function of the gene is lost (dashed triangle) and the truncated protein has a deleterious effect (red triangle). This leads to a greater divergence between the phenotype and the target (orange area on the phenotype). Bottom: The deletion removes five genes. The functions of two of them can be seen in the box (dotted triangles). This results in a large discrepancy between the phenotype and the target (orange area on the phenotype).

All individuals are replaced at each generation following a spatialized Wright-Fisher model. The number of descendants of each individual depends on its fitness difference with its neighbors. At each reproduction event, point mutations or genomic rearrangements can occur (Figure 7C). They create diversity in the genomes, hence in the phenotypes, and allow the genome size and structure to change. These changes can be neutral or not, depending on whether mutations alter coding and/or non-coding sequences. These changes do not have a predefined effect on the fitness of the offsprings as their genomes will be decoded thereafter, thus the model do not impose an *a priori* genome structure and allows us to study the evolution of genome architecture in various experimental conditions.

The mutation rate (in *bp*^*−*1^) is set for each type of mutation independently. When all mutation rates are equal, there is in an equal probability of losing or gaining base pairs. The size distribution of InDels is uniform in [1, 6], and the size distribution of large deletions and duplications is uniform in [1, *L*] (with *L* the genome length).

### 4.2 Experimental design

#### 4.2.1 Wild Types

In order to observe changes in genome architecture induced by changes in the population size and/or mutation rates, we begin our experiments from pre-evolved organisms, which are called “Wild Types” (WT). Having already evolved for millions of generations under constant conditions, WTs are very stable in genome structure and well adapted to their environment (although the fitness never stops increasing). 5 different WTs were used for our experiments, all having evolved for 10 millions generations at the basal conditions of *N*_0_ = 1, 024 individuals and a mutation rate of *µ*_0_ = 10^*−*6^ mutations per base pair per generation for each type of mutations (point mutations, small insertions, small deletions, inversions, duplications, large deletions and translocations). Importantly, in this experiment, all types of mutations are equally probable: there is no mutational bias towards the insertion or deletion of base pairs. Bacterial populations are very large and cannot be directly modeled owing to computational load. We hence limit the population sizes in our experiments, but compensated by increasing the mutation rates such that the *N × µ* parameter is of the same order of magnitude as for real bacterial populations. Finally, in order to limit the effect of drift, we used a selection strength *k* = 1, 000, which is relatively high and guarantees an efficient selection. The fitnesses and genome structures of the WTs are listed in Table 1.

#### 4.2.2 Experimental conditions

A range of population sizes increases or decreases and mutations rates increases, as well as some combinations of both, are tested. All conditions are listed in Table 2 below. For each combination of conditions, 10 replications of each of the 5 WTs are run. Initial populations are always clonal : all individuals are identical to the specific WT used for the run.

**Table 2:** Experimental conditions tested. The control condition is in bold. Note that, as the simulations take place on a squared grid, population sizes could not be exactly divided or multiplied by 2.

| Population size | Mutation rate<br>(per base pair,<br>per mutation type) | $N \times \mu$<br>product |
| --- | --- | --- |
| 64 ($N_0/16$) | $10^{-6}$ ($\mu_0$) | $1/16 N_0 \times \mu_0$ |
| 256 ($N_0/4$) | $10^{-6}$ ($\mu_0$) | $1/4 N_0 \times \mu_0$ |
| **1024 ($N_0$)** | **$10^{-6}$ ($\mu_0$)** | $N_0 \times \mu_0$ |
| 529 ($\approx N_0/2$) | $2 \times 10^{-6}$ ($2 \times \mu_0$) | $\approx N_0 \times \mu_0$ |
| 256 ($N_0/4$) | $4 \times 10^{-6}$ ($4 \times \mu_0$) | $N_0 \times \mu_0$ |
| 64 ($N_0/16$) | $16 \times 10^{-6}$ ($16 \times \mu_0$) | $N_0 \times \mu_0$ |
| 2,025 ($\approx 2 \times N_0$) | $2 \times 10^{-6}$ ($2 \times \mu_0$) | $\approx 4 N_0 \times \mu_0$ |
| 4,096 ($4 \times N_0$) | $10^{-6}$ ($\mu_0$) | $4 N_0 \times \mu_0$ |
| 1,024 ($N_0$) | $4 \times 10^{-6}$ ($4 \times \mu_0$) | $4 N_0 \times \mu_0$ |
| 4,096 ($4 \times N_0$) | $4 \times 10^{-6}$ ($4 \times \mu_0$) | $16 N_0 \times \mu_0$ |
| 16,384 ($16 \times N_0$) | $10^{-6}$ ($\mu_0$) | $16 N_0 \times \mu_0$ |
| 1,024 ($N_0$) | $16 \times 10^{-6}$ ($16 \times \mu_0$) | $16 N_0 \times \mu_0$ |
| 16,384 ($16 \times N_0$) | $16 \times 10^{-6}$ ($16 \times \mu_0$) | $16 N_0 \times \mu_0$ |

#### 4.2.3 Data analyses

To analyze the simulations, we reconstruct the ancestral lineages of the final populations. To this end, simulations are run for 2, 100, 000 generations, and we identify all the ancestors of a random individual of the final population. We then study the data from generation 0 to generation 2, 000, 000 and ignore the last 100, 000 to ensure that the final population has coalesced and that we study the lineage of the whole final population.

On this lineage, we retrieve the fitness, coding and non-coding genome size at each generation, as well as the replicative robustness every 1, 000 generations. The replicative robustness is measured as the proportion of the offspring of an individual which has the exact same fitness as its parent, *i.e*. that underwent no mutation at all, or only purely neutral mutations. To estimate replicative robustness for a given individual of the lineage, we generate 10, 000 offsprings and compare them to their parent.

To compare experimental conditions, we retrieve the individual at generation 2, 000, 000 in each lineage. This individual is the common ancestor of the final population (at generation 2, 100, 000), thus ensuring that its genome structure has been conserved by evolution. A visualization of the temporal lineage data (fitness, coding fraction and total, coding and non-coding genome sizes) for the 50 replicas of each experimental condition is provided in the supplementary material S3 (Figures S3 to S15).

#### 4.2.4 Effect of mutational biases

As it is often assumed that mutational biases – towards deletions for bacteria and towards insertions for eukaryotes – are very important for genome size evolution (Petrov, 2002), we also tried to confront our experiments to the impact of mutational biases. We tested four mutational biases : twice as many large deletions than duplications, twice as many small deletions than small insertions, twice as many duplications than large deletions, twice as many small insertions than small deletions. In all cases, the sum of all mutation rates is conserved such that the overall mutational pressure is the same as in the previous experiments.

For each mutational condition, 5 Wild-Types evolved for 10, 000, 000 generations. Then, the median-sized WT of each mutational condition was extracted and confronted to either an increase or decrease in population size (4 *× N*_0_, *N*_0_*/*4), or an increase in all mutation rates proportionally (4 *× µ*_0_ – note that, in case of bias, *µ*_0_ may be different for the different types of mutation) for 2, 100, 000 generations. By extracting the ancestor of the lineage at generation 2, 000, 000, we could compare these experiments to the control conditions (where the population size and mutation rates remained stable for 2, 100, 000 generations).

#### 4.2.5 Data availability

The code of Aevol is available on gitlab at https://gitlab.inria.fr/aevol/aevol. WTs sequences to reproduce the experiments, as well as the full lineages data and robustness data, are available on Zenodo : https://zenodo.org/doi/10.5281/zenodo.10669479.

## 5 Acknowledgments

The authors acknowledge the support of the French Agence Nationale de la Recherche (ANR), under grant ANR-20-CE02-0008 (NeGA project). J.L., G.B. and N.L. would like to thank the Rhône-Alpes Institute for Complex Systems (IXXI) for funding. All authors thank the Grid’5000 testbed, supported by a scientific interest group hosted by Inria and including CNRS, RENATER and several Universities as well as other organizations (see https://www.grid5000.fr), for computational support. The authors would like to thank Laurent Duret and David P. Parsons for fruitful comments on the manuscript.

