## Supplementary Materials for "Genome streamlining: effect of mutation rate and population size on genome size reduction"

#### S1 Effective population size in a model with local competition.

Based on the work of Zhang et al. 2014, we computed the theoretical effective population size in Aevol, when the reproduction is limited to the direct neighborhood of the organism (9 possibilities). It appears that, approximately, $Ne \propto N\log(N)$. The difference is however relatively small for low $N$ values.

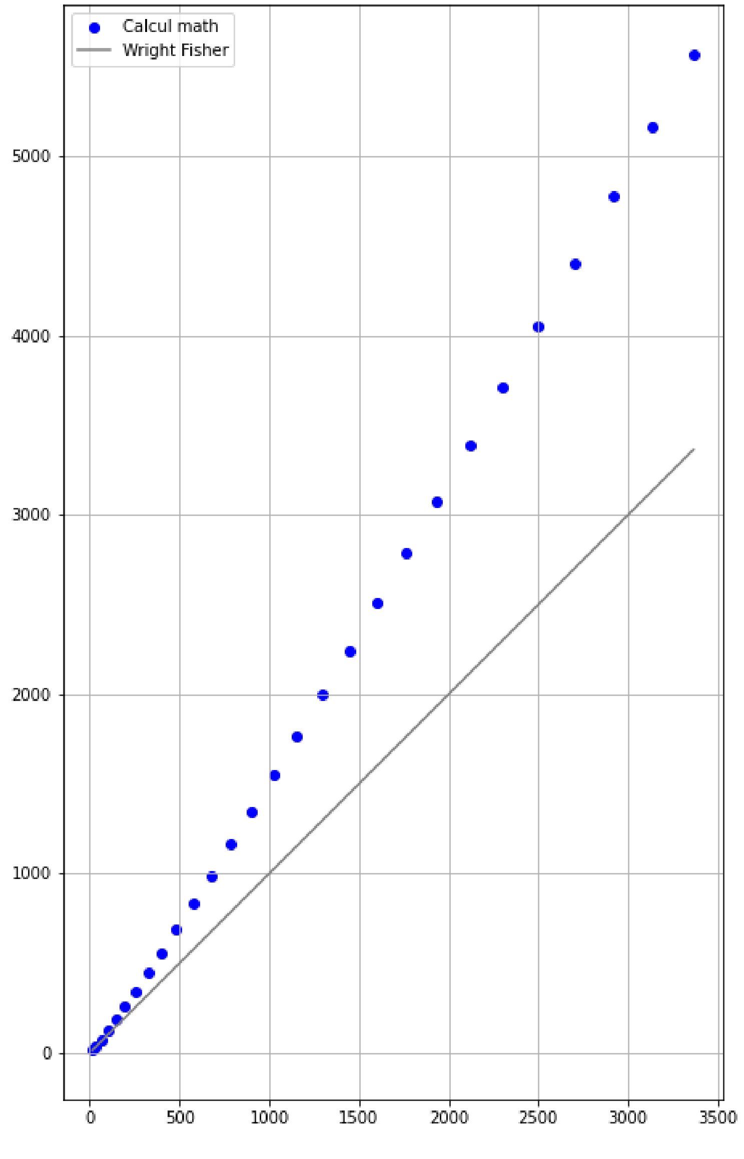


Figure S1: $Ne$ as a function of $N$ for a Wright-Fischer model and for a model with local reproduction. This supposes the absence of selective sweep.

Note that, to divide the effective population size by 16 when starting from $N = 1024$, one should use $N = 81$ instead of $N = 64$. Some simulations tested with $N = 81$ and $\mu = 1.6 \times 10^{-6}$ do show that in these conditions the coding percentage goes closer to the initial value of 0.68% (*data not shown*).

#### S2 Results with a mutational bias in the InDels.

To test the interaction between mutational biases and changes in population size or mutation rate, we also evolved 5 Wild-Types with an insertion bias in the InDels distribution (twice more insertions than deletions), or a deletion bias (twice more deletions than insertions), keeping the total mutation rate constant. Similarly to what is described in the main text with the bias in the larger events, the genome sizes of our Wild-Types are different from the one without a mutational bias: $28,941$ and a coding fraction of $0.37$ with the insertion bias *VS* $8,925$ and $0.98$ with a deletion bias.

When submitted to a change in population size or mutation rate, the median WT of both these experiments react quite similarly to what is predicted by our model (see Fig. S2) —although the loss of non-coding bases is quite limited in the deletion-bias experiment since the coding proportion is already so high.

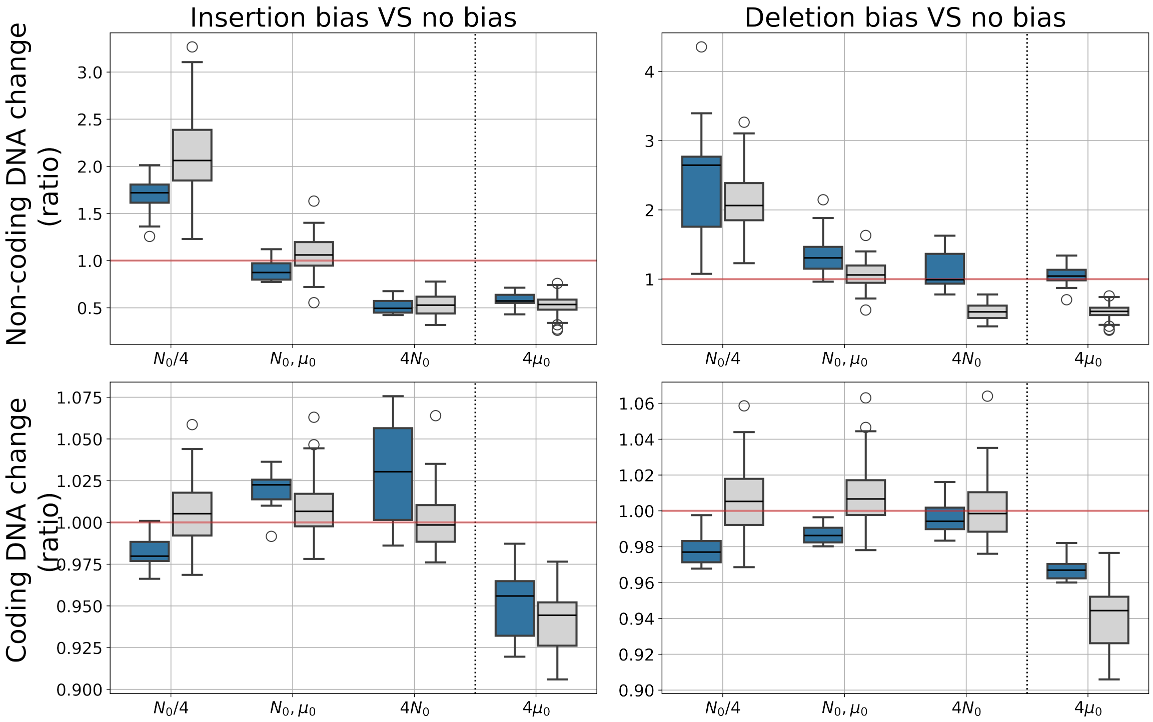


Figure S2: Change in coding and non-coding genome sizes in reaction to changes in $N$ or $\mu$ for the different mutational biases. Blue boxes show the results with a mutational bias (left: insertion bias in InDels, right: deletion bias in InDels), and gray boxes show the results without mutational bias. Depicted values are the ratio of the coding/non-coding size at the final generation over the value at generation 0.

### S3 Temporal data for all tested conditions.

For each of the conditions tested (see Table 1 of the main document, section 4.2.2), we provide the temporal data of fitness, total amount of DNA, coding and non-coding sizes as well as the coding fraction for all 50 repetitions. Curves are colored by WT used to start the simulation (blue: WT1, orange: WT2, green: WT3, purple: WT4, brown: WT5). Individual simulations are depicted as shallow lines, and the thick curves show the average value for the Wild-Type.

#### 3.1 Control: $\mu = 10^{-6} = \mu_0, \quad N = 1024 = N_0, \quad N \times \mu = N_0 \times \mu_0$

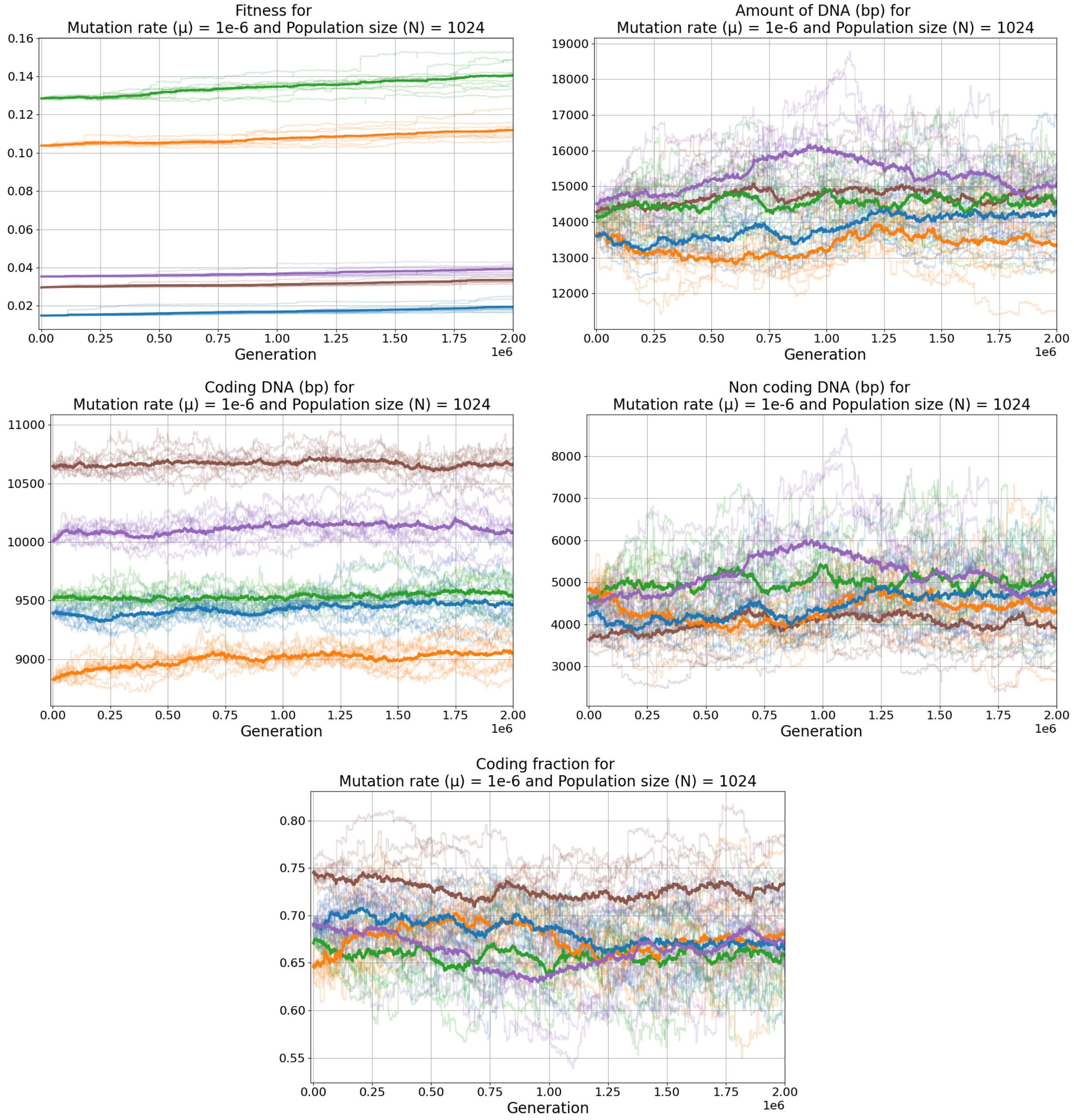


Figure S3: Temporal data for $\mu = 10^{-6}$, $N = 1024$. From left to right, top to bottom : Fitness, total amount of DNA, coding genome size, non-coding genome size and coding fraction.

##### 3.2 $\mu = 10^{-6} = \mu_0, \quad N = 64 = N_0/16, \quad N \times \mu = 1/16 N_0 \times \mu_0$

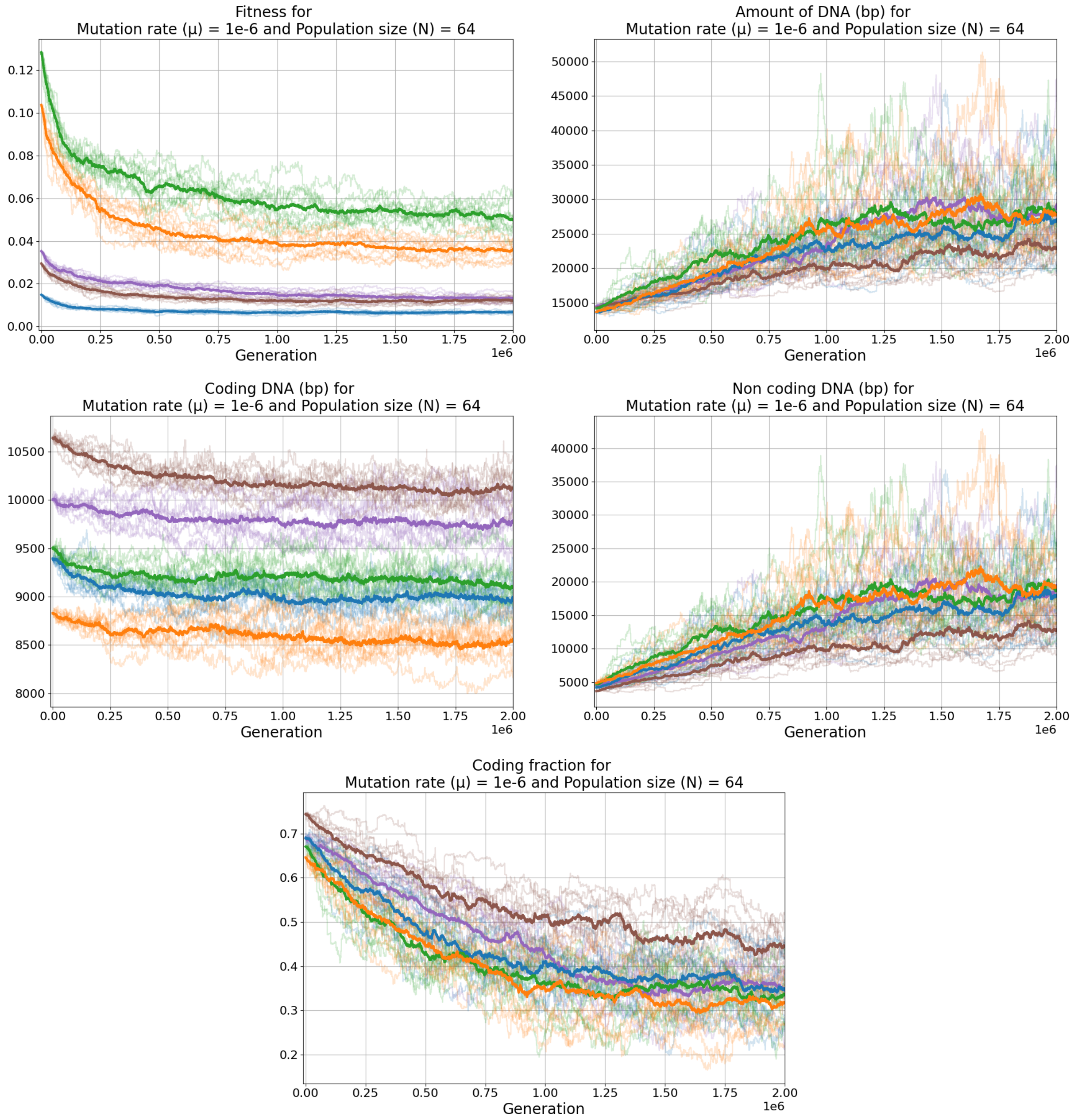


Figure S4: Temporal data for $\mu = 10^{-6}$, $N = 64$. From left to right, top to bottom : Fitness, total amount of DNA, coding genome size, non-coding genome size and coding fraction.

#### 3.3 $\mu = \mu_0 = 10^{-6}$, $N = N_0/4 = 256$, $N \times \mu = 1/4 N_0 \times \mu_0$

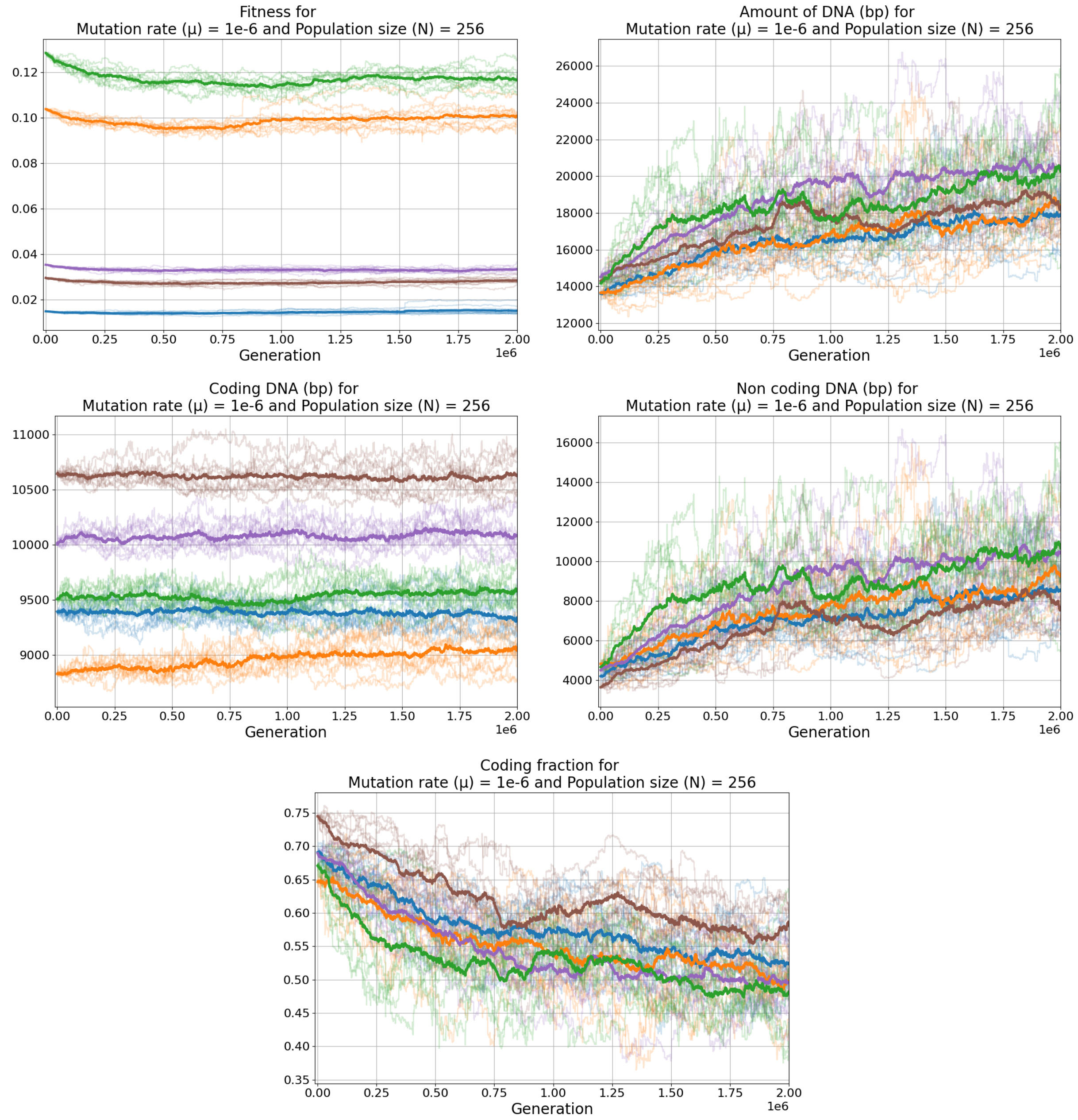


Figure S5: Temporal data for $\mu = 10^{-6}$, $N = 256$. From left to right, top to bottom : Fitness, total amount of DNA, coding genome size, non-coding genome size and coding fraction.

#### 3.4 $\mu = 10^{-6} = \mu_0, \quad N = 4096 = 4N_0, \quad N \times \mu = 4N_0 \times \mu_0$

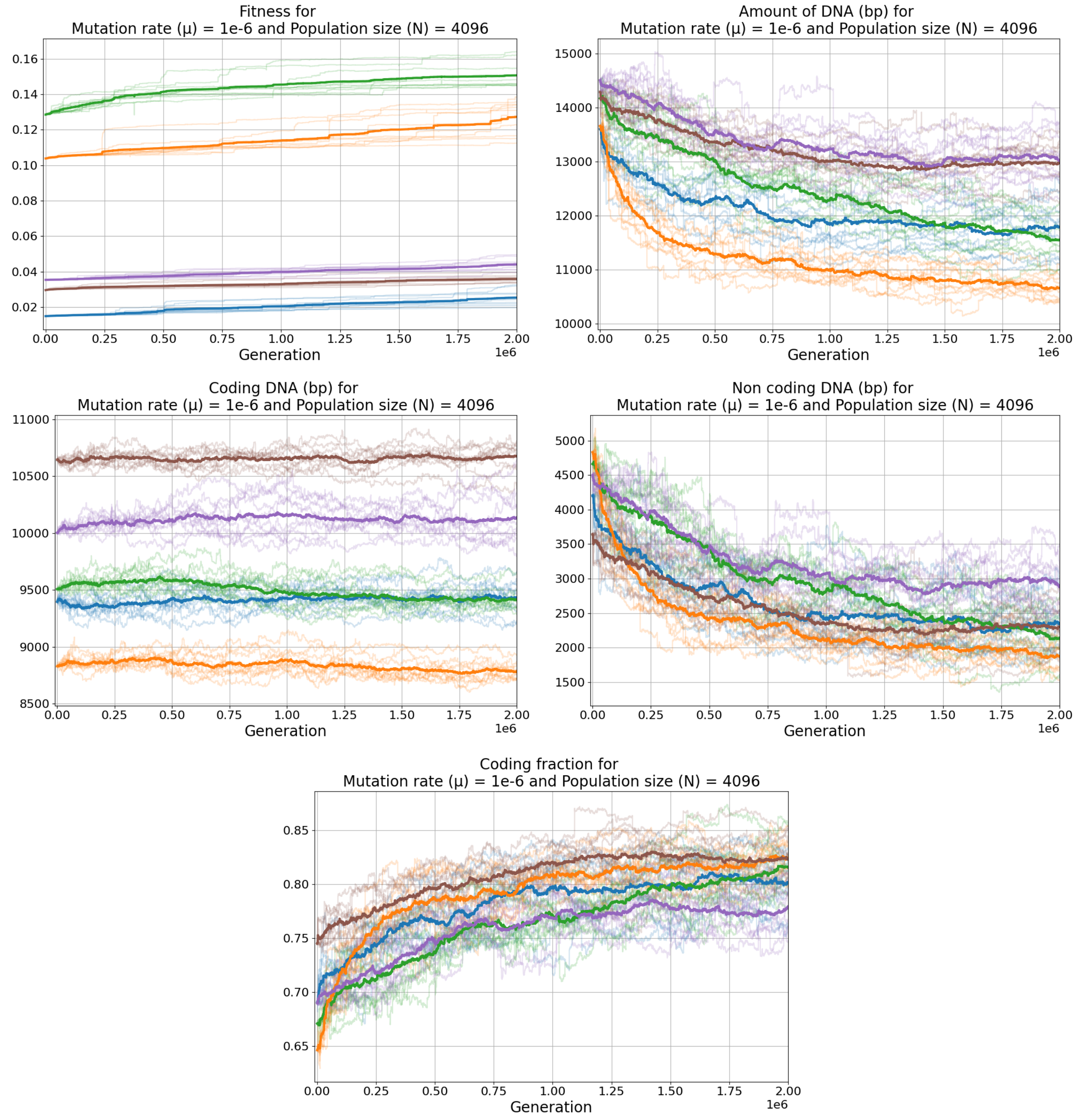


Figure S6: Temporal data for $\mu = 10^{-6}$, $N = 4096$. From left to right, top to bottom : Fitness, total amount of DNA, coding genome size, non-coding genome size and coding fraction.

#### 3.5 $\mu = \mu_0 = 10^{-6}$, $N = 16N_0 = 16384$, $N \times \mu = 16N_0 \times \mu_0$

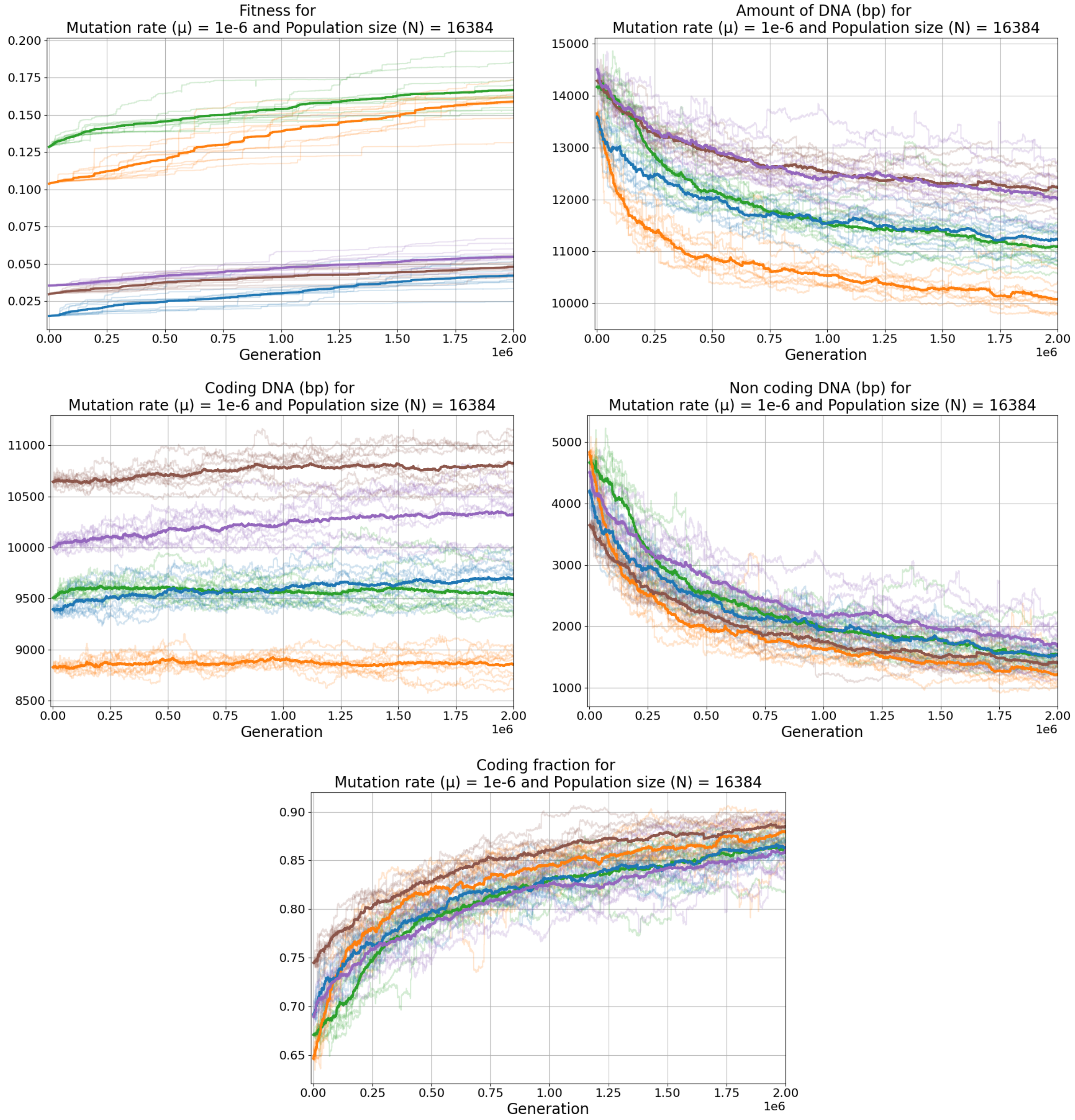


Figure S7: Temporal data for $\mu = 10^{-6}$, $N = 16384$. From left to right, top to bottom : Fitness, total amount of DNA, coding genome size, non-coding genome size and coding fraction.

#### 3.6 $\mu = 2 \times 10^{-6} = 2\mu_0, \quad N = 529 \approx N_0/2, \quad N \times \mu \approx N_0 \times \mu_0$

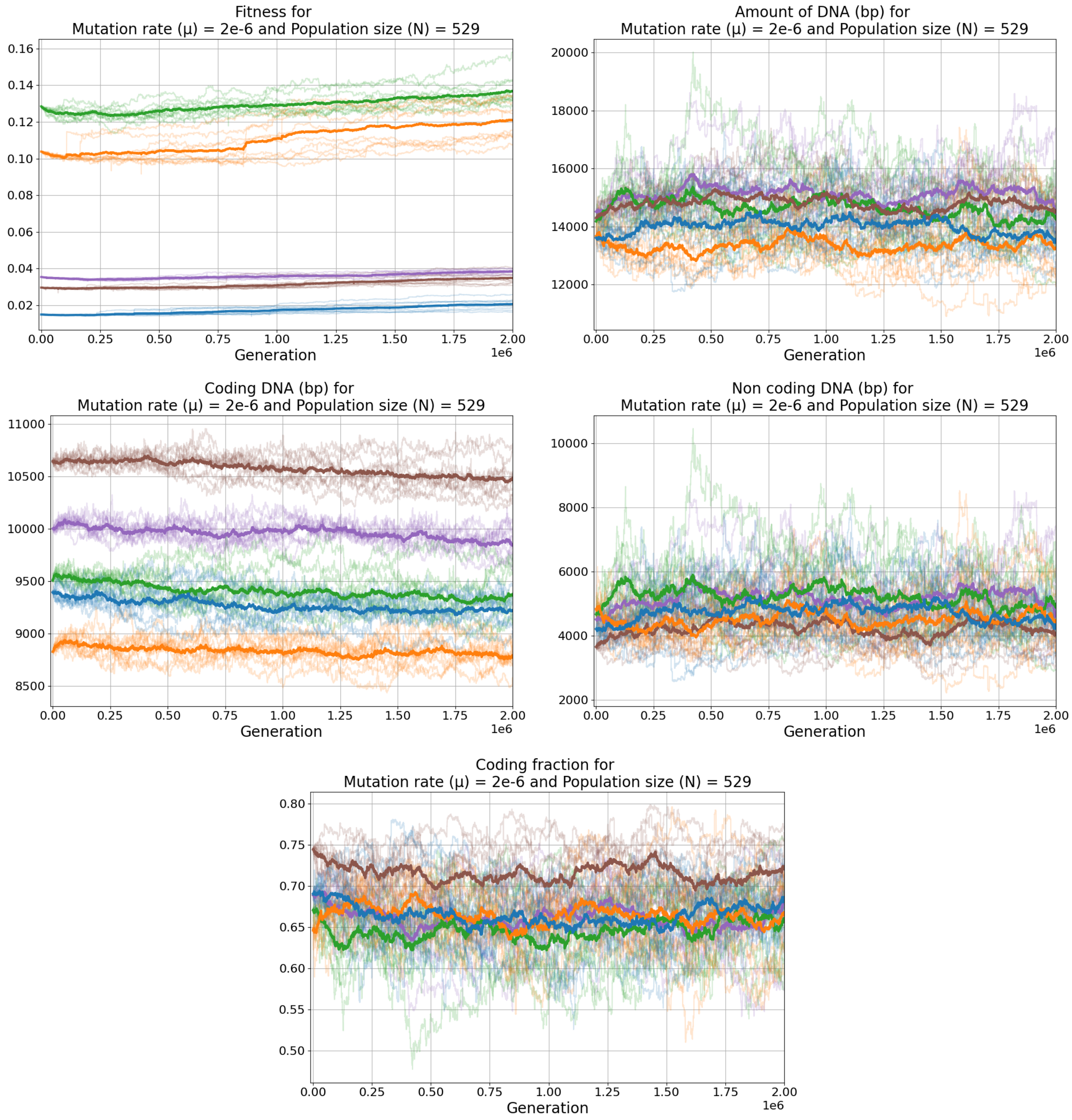


Figure S8: Temporal data for $\mu = 2 \times 10^{-6}$, $N = 529$. From left to right, top to bottom : Fitness, total amount of DNA, coding genome size, non-coding genome size and coding fraction.

##### 3.7 $\mu = 2 \times 10^{-6} = 2\mu_0, \quad N = 2025 \approx 2N_0, \quad N \times \mu \approx 4N_0 \times \mu_0$

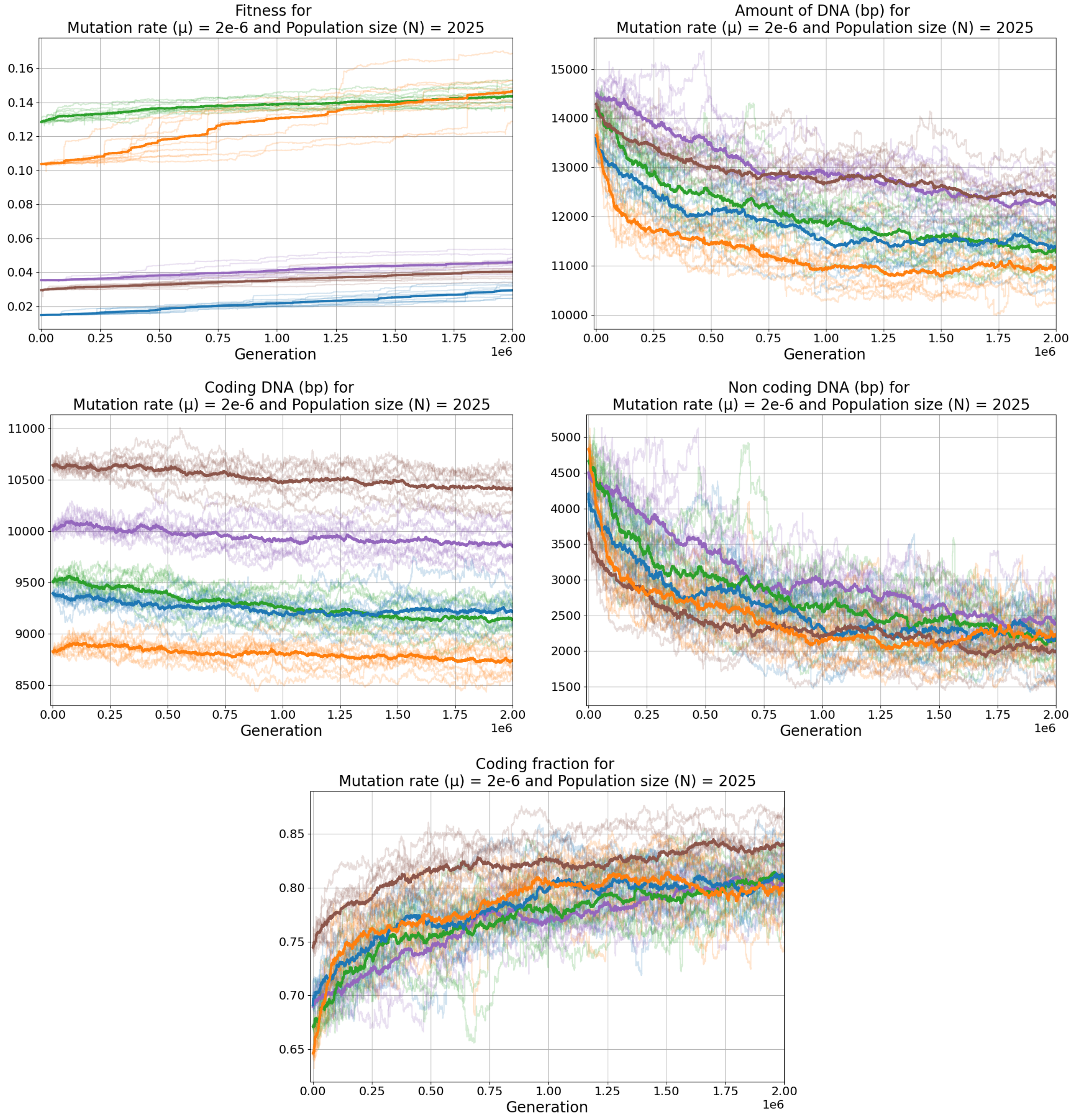


Figure S9: Temporal data for $\mu = 2 \times 10^{-6}$, $N = 2025$. From left to right, top to bottom : Fitness, total amount of DNA, coding genome size, non-coding genome size and coding fraction.

#### 3.8 $\mu = 4 \times 10^{-6} = 4\mu_0, \quad N = 256 = N_0/4, \quad N \times \mu = N_0 \times \mu_0$

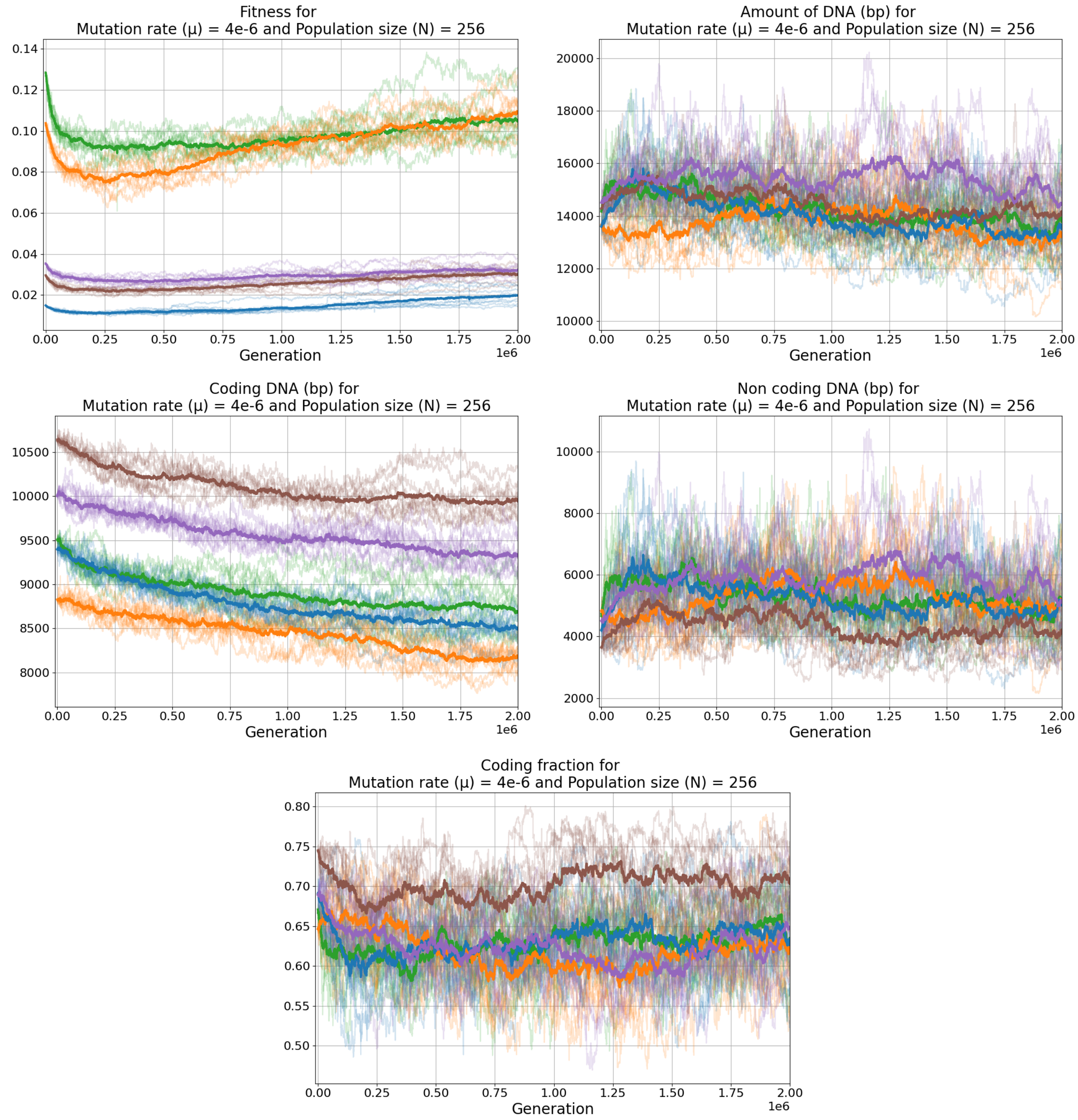


Figure S10: Temporal data for $\mu = 4 \times 10^{-6}$, $N = 256$. From left to right, top to bottom : Fitness, total amount of DNA, coding genome size, non-coding genome size and coding fraction.

#### 3.9 $\mu = 4 \times 10^{-6} = 4\mu_0, \quad N = 1024 = N_0, \quad N \times \mu = 4N_0 \times \mu_0$

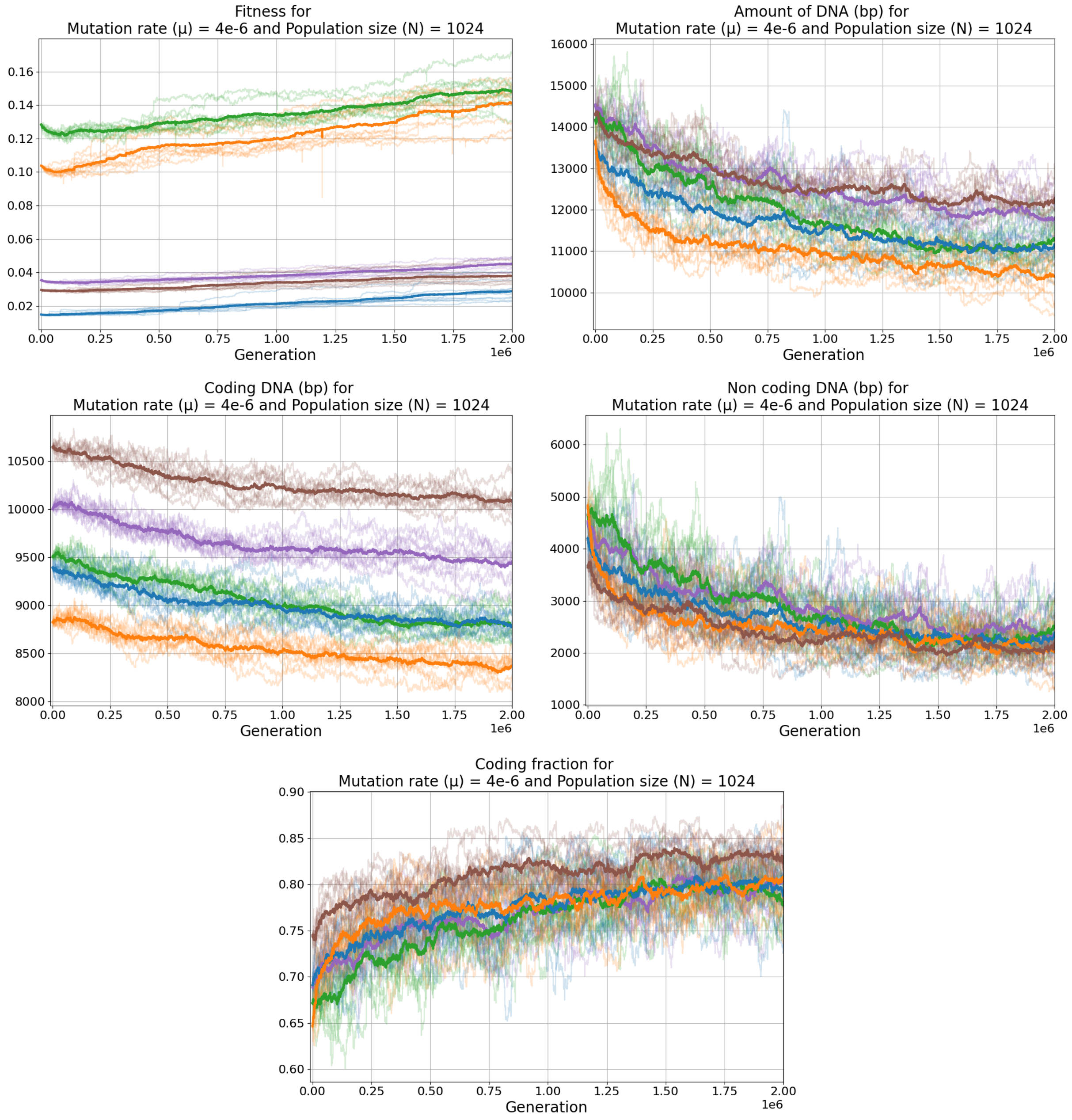


Figure S11: Temporal data for $\mu = 4 \times 10^{-6}$, $N = 1024$. From left to right, top to bottom : Fitness, total amount of DNA, coding genome size, non-coding genome size and coding fraction.

#### 3.10 $\mu = 4 \times 10^{-6} = 4\mu_0, \quad N = 4096 = 4N_0, \quad N \times \mu = 16N_0 \times \mu_0$

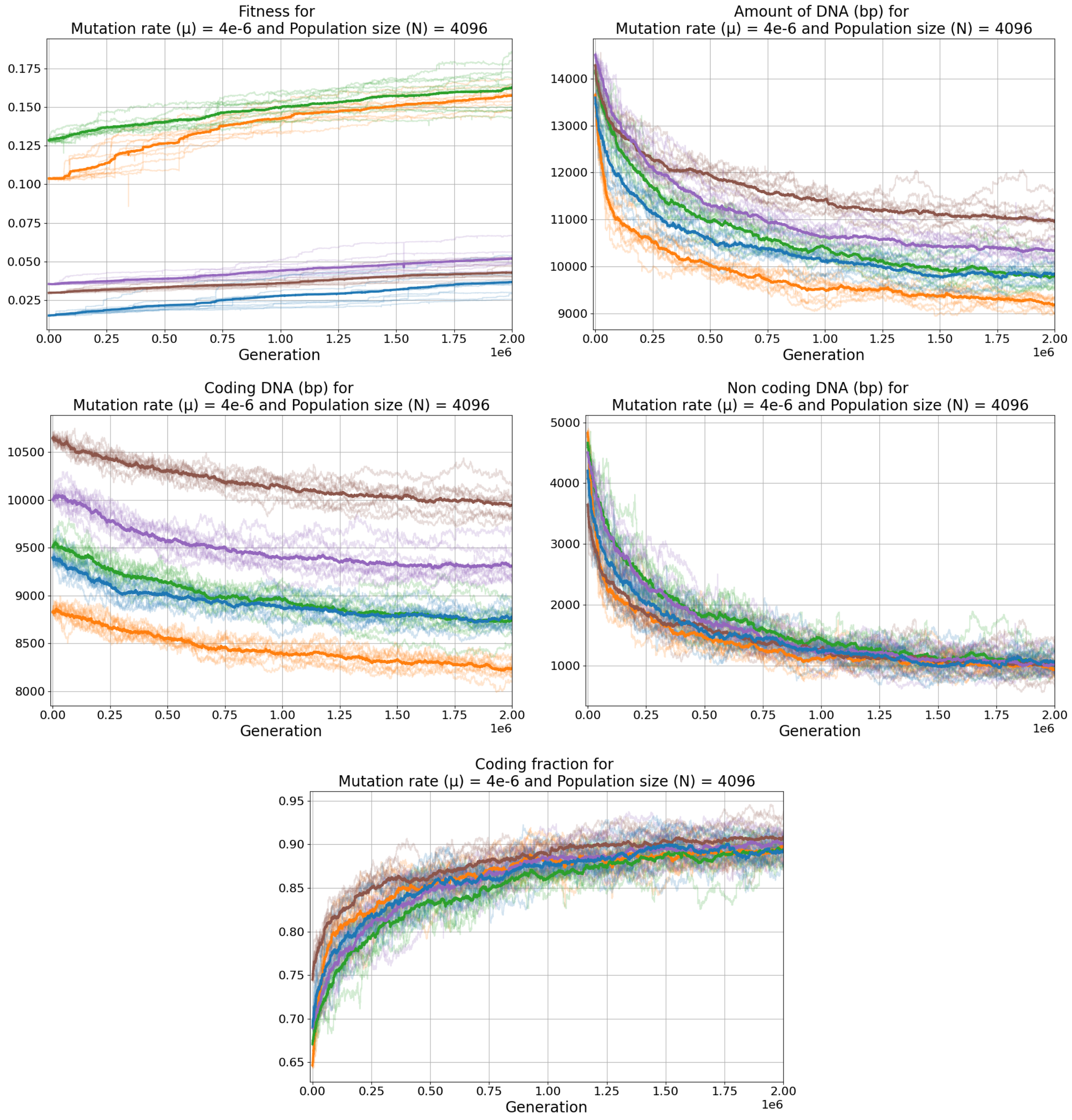


Figure S12: Temporal data for $\mu = 4 \times 10^{-6}$, $N = 4096$. From left to right, top to bottom : Fitness, total amount of DNA, coding genome size, non-coding genome size and coding fraction.

#### 3.11 $\mu = 1.6 \times 10^{-5} = 16\mu_0$, $N = 64 = N_0/16$, $N \times \mu = N_0 \times \mu_0$

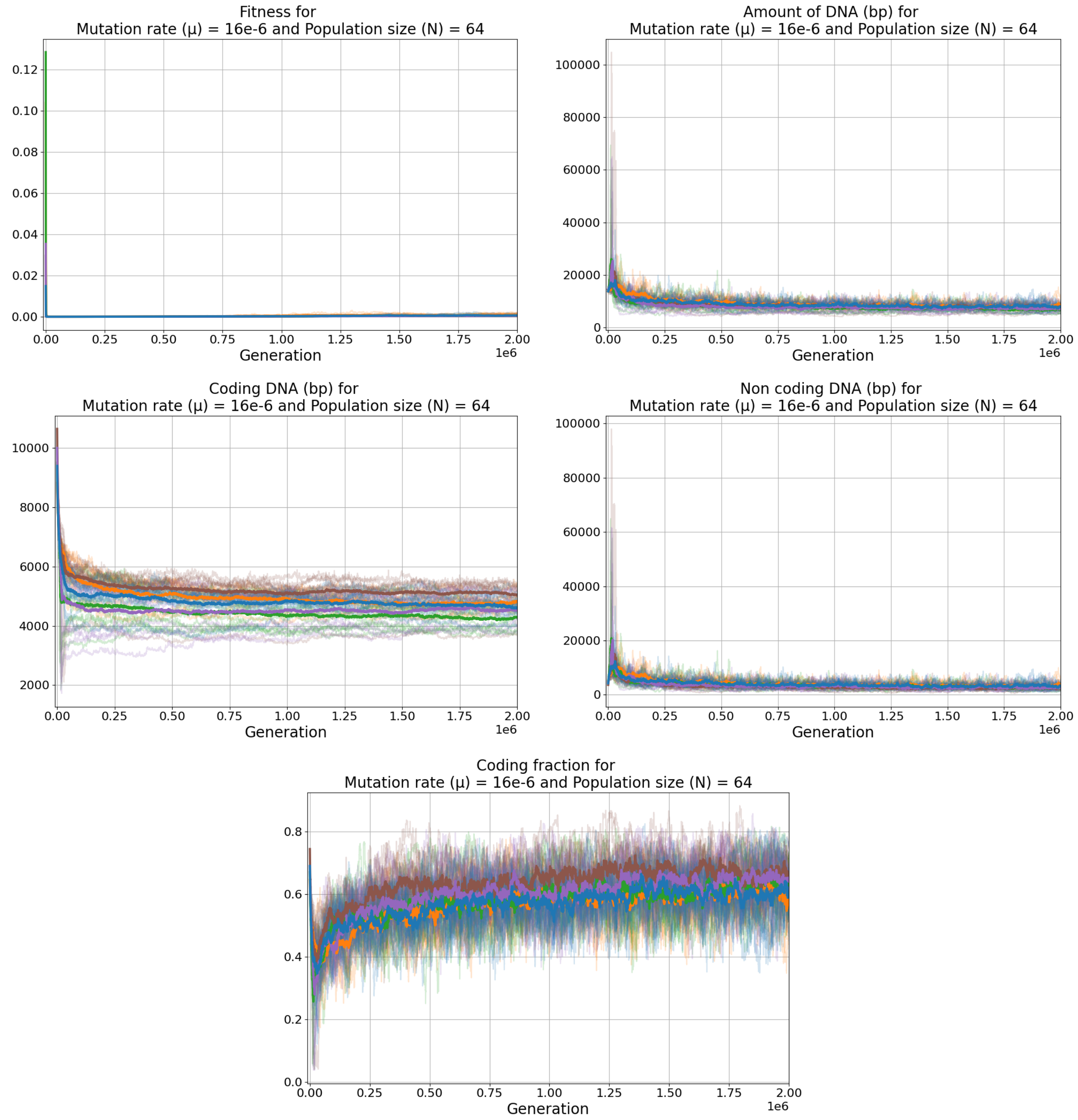


Figure S13: Temporal data for $\mu = 1.6 \times 10^{-5}$, $N = 64$. From left to right, top to bottom : Fitness, total amount of DNA, coding genome size, non-coding genome size and coding fraction.

#### 3.12 $\mu = 1.6 \times 10^{-5} = 16\mu_0, \quad N = 1024 = N_0, \quad N \times \mu = 16N_0 \times \mu_0$

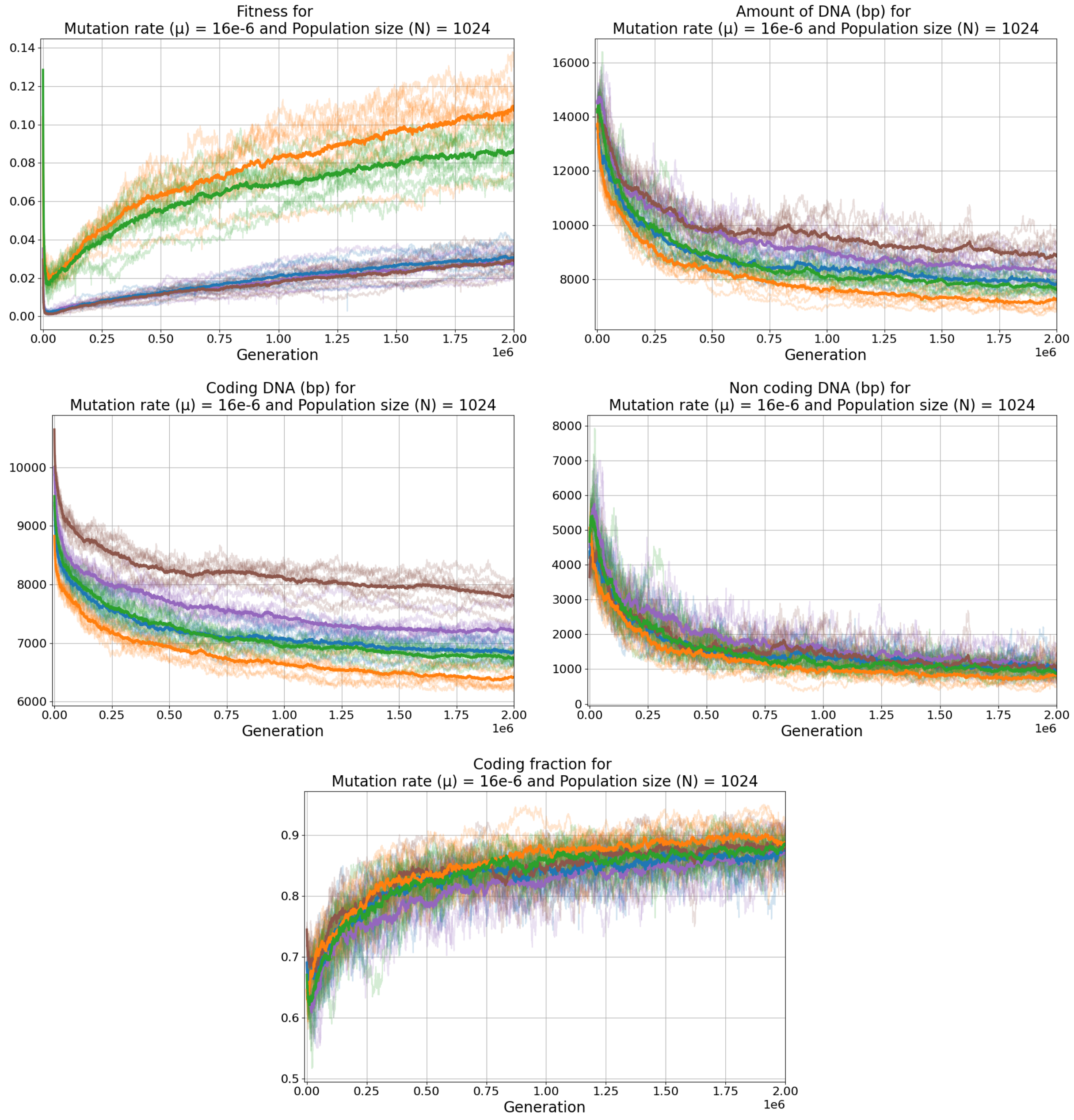


Figure S14: Temporal data for $\mu = 1.6 \times 10^{-5}$, $N = 1024$. From left to right, top to bottom : Fitness, total amount of DNA, coding genome size, non-coding genome size and coding fraction.

#### 3.13 $\mu = 1.6 \times 10^{-5} = 16\mu_0, \quad N = 16,384 = 16N_0, \quad N \times \mu = 256N_0 \times \mu_0$

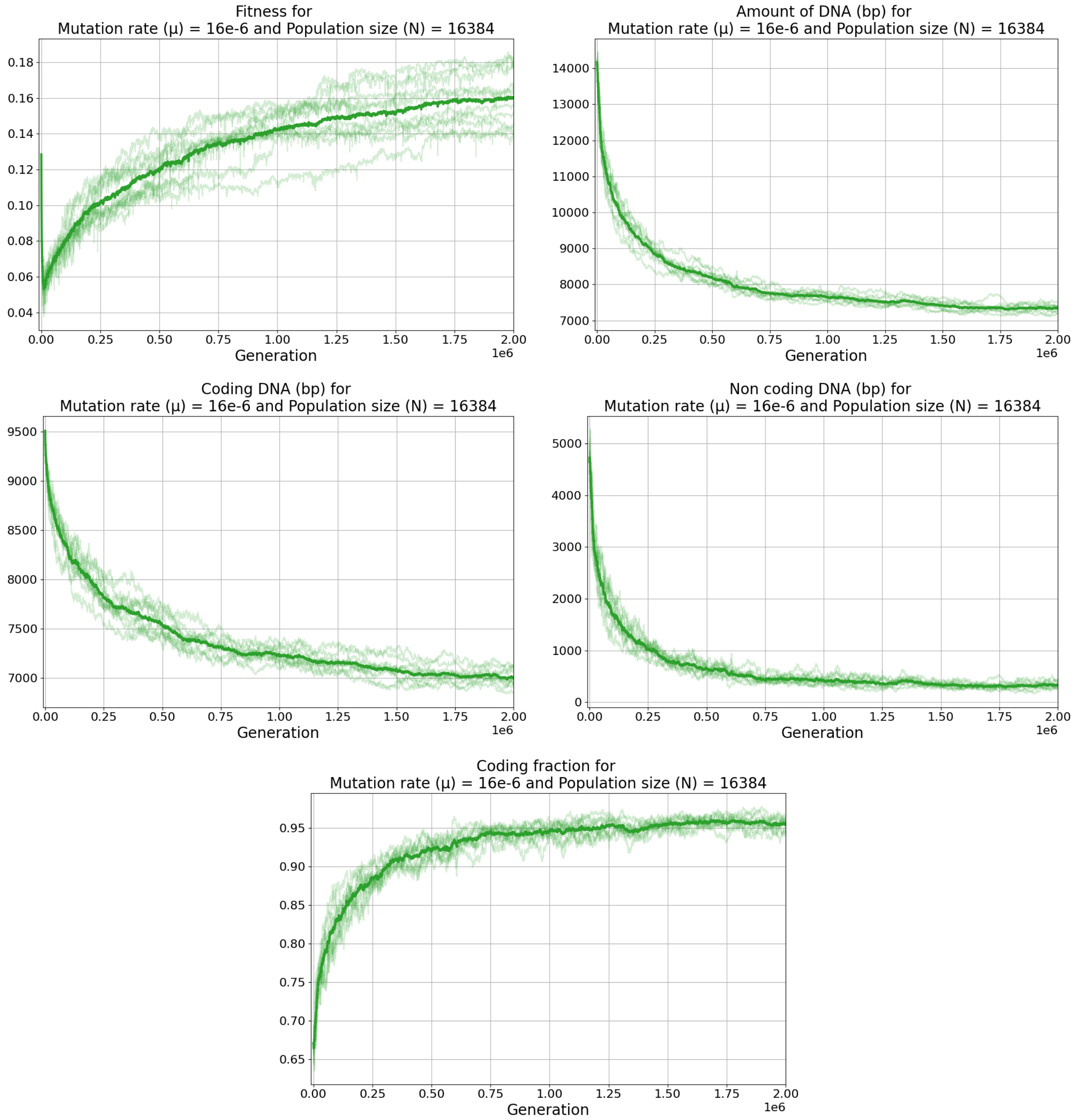


Figure S15: Temporal data for $\mu = 1.6 \times 10^{-5}$, $N = 16,384$. From left to right, top to bottom : Fitness, total amount of DNA, coding genome size, non-coding genome size and coding fraction.
